# Global architecture of the nucleus in single cells by DNA seqFISH+ and multiplexed immunofluorescence

**DOI:** 10.1101/2020.11.29.403055

**Authors:** Yodai Takei, Jina Yun, Noah Ollikainen, Shiwei Zheng, Nico Pierson, Jonathan White, Sheel Shah, Julian Thomassie, Chee-Huat Linus Eng, Mitchell Guttman, Guo-cheng Yuan, Long Cai

## Abstract

Identifying the relationships between chromosome structures, chromatin states, and gene expression is an overarching goal of nuclear organization studies. Because individual cells are highly variable at all three levels, it is essential to map all three modalities in the same single cell, a task that has been difficult to accomplish with existing tools. Here, we report the direct super-resolution imaging of over 3,660 chromosomal loci in single mouse embryonic stem cells (mESCs) by DNA seqFISH+, along with 17 chromatin marks by sequential immunofluorescence (IF) and the expression profile of 70 RNAs, in the same cells. We discovered that the nucleus is separated into zones defined by distinct combinatorial chromatin marks. DNA loci and nascent transcripts are enriched at the interfaces between specific nuclear zones, and the level of gene expression correlates with an association between active or nuclear speckle zones. Our analysis also uncovered several distinct mESCs subpopulations with characteristic combinatorial chromatin states that extend beyond known transcriptional states, suggesting that the metastable states of mESCs are more complex than previously appreciated. Using clonal analysis, we show that the global levels of some chromatin marks, such as H3K27me3 and macroH2A1 (mH2A1), are heritable over at least 3-4 generations, whereas other marks fluctuate on a faster time scale. The long-lived chromatin states may represent “hidden variables” that explain the observed functional heterogeneity in differentiation decisions in single mESCs. Our integrated spatial genomics approach can be used to further explore the existence and biological relevance of molecular heterogeneity within cell populations in diverse biological systems.

## Main

Currently, the main approaches to examine nuclear organization are 1) sequencing-based genomics, which measures contacts between DNA loci^1–4^ and between DNA and nuclear bodies^5–10^, and 2) microscopy-based imaging of chromosomes in single cells, conventionally by multicolor DNA fluorescence in situ hybridization (DNA FISH)^11,12^. Genomics approaches have been powerful in mapping global contacts between chromosomes and have been scaled down to the single cell level^13–19^. However, reconstructing 3D structures from the measured pairwise contacts relies on computational models and assumptions. In addition, it is difficult to integrate multiple modalities of measurements in the same cells. On the other hand, recent imaging methods^20–24^ that walk along the chromosome by sequential DNA FISH or oligo-stochastic optical reconstruction microscopy (oligo-STORM), provide a direct view of chromosomes in single cells. These studies have confirmed genomic features such as topological associating domains found by Hi-C^20,21,23,24^ and showed that chromosome organization is highly heterogeneous at the single cell level^21,24,25^.

### DNA seqFISH+ images global chromosome conformation in single cell

Integrating multiple modalities in imaging-based approaches could reveal new insights into nuclear organization beyond those obtained by sequencing-based methods. We previously demonstrated DNA sequential fluorescence in situ hybridization (DNA seqFISH) with 12 subtelomeric regions in single nuclei^26^. We recently showed that over 10,000 genes can be multiplexed at the transcription active sites in single cells with intron seqFISH^27^, and that a deterministic super-resolution version called RNA seqFISH+ can overcome the optical density issues in single cells to profile mRNAs at the transcriptome scale^28^.

Building upon these previous work, we now developed DNA seqFISH+ to target over 3,660 loci in single mouse embryonic stem cells (mESCs) (Fig. 1a, Extended Data Fig. 1a, Supplementary Table 1). Briefly, we used 16 rounds of hybridization in each fluorescent channel to generate a super-resolved image that is repeated for a total of 5 times to barcode up to 2,048 barcodes with 2 round error correction (Fig. 1b, c, Extended Data Fig. 1b, Supplementary Table 2). The barcoding is performed within a single fluorescent channel to avoid chromatic aberration during barcode calling steps. In the first fluorescent channel, 1,267 loci were selected approximately 2 megabases (Mb) apart to cover the whole genome uniformly, and include gene-poor regions. The second fluorescent channel targeted 1,193 loci at the 5’ end of genes, including those related to pluripotency and differentiation of mESCs. Together these two channels label 2,460 loci spaced approximately 1 Mb apart across the whole genome. At each locus targeted, we used up to 200 probes within the 25kb region and imaged individual loci as diffraction limited spots. The third fluorescent channel targets 60 consecutive loci at 25 kilobase (kb) resolution on each of the 20 chromosomes for an additional 1,200 loci (Fig. 1b, d, Extended Data Fig. 1c, Supplementary Table 2). In each of the first 60 rounds of hybridizations in this channel, 20 loci (one locus on each chromosome) were imaged simultaneously. The last 20 rounds of hybridization were used to sequentially “paint” individual chromosomes and assign a chromosome identity to each of the distinct regions in the 20 chromosomes. Overall, 80 rounds of hybridization are used for the DNA seqFISH+ experiment (Extended Data Fig. 2a-c). This approach allowed us to examine nuclei at both 1 Mb resolution globally for the entire genome, and 25 kb resolution for 20 distinct regions that are at least 1.5 Mb in size.

**Figure 1.**
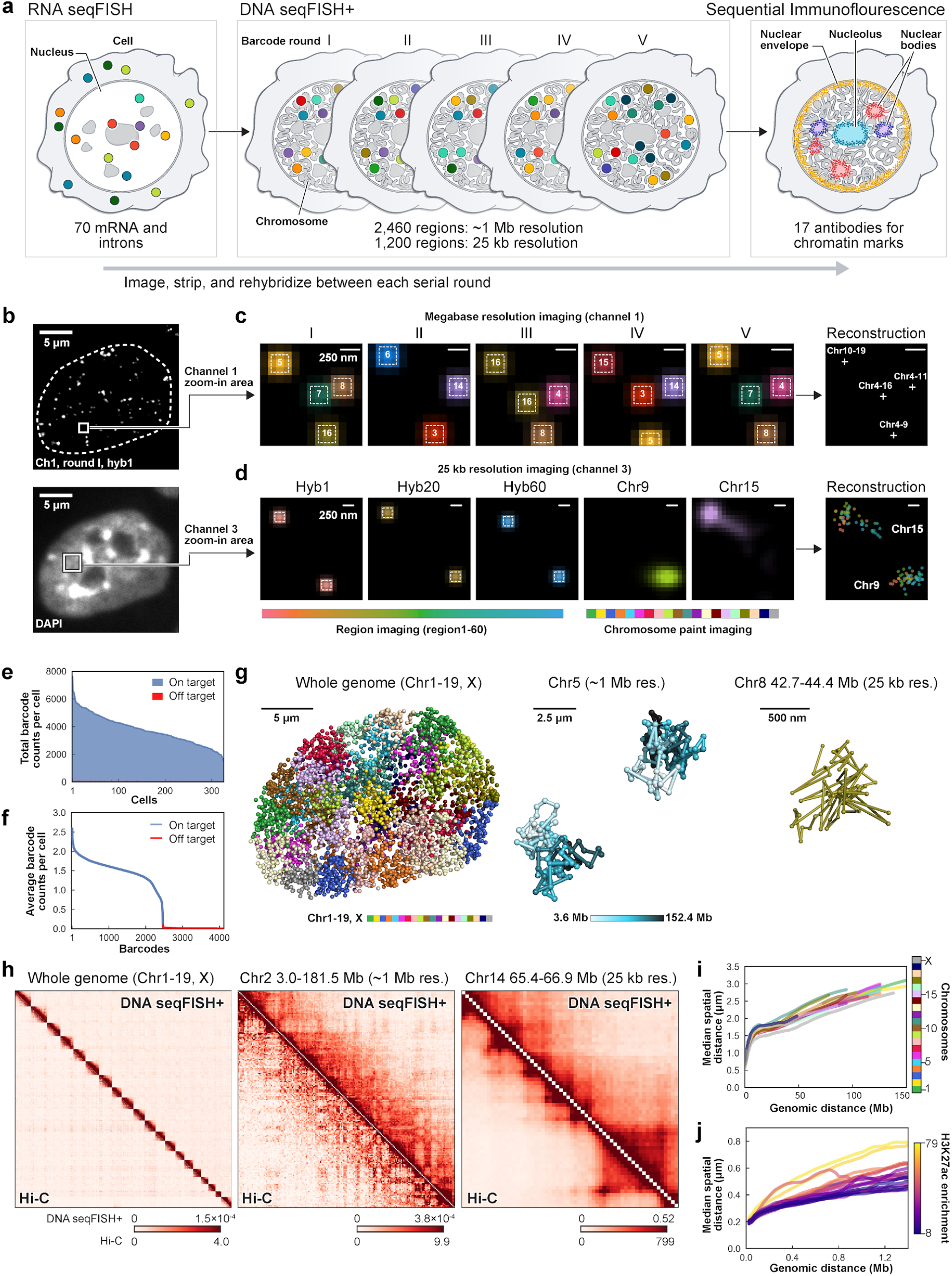
DNA seqFISH+ enables large-scale super-resolution imaging of chromosomes. a, Schematic for DNA seqFISH+ combined with RNA seqFISH and sequential IF. 70 RNA species targeting mRNA and nascent RNA are first imaged by RNA seqFISH. Then, a total of 3,660 regions, consisting of 2,460 regions with approximately 1 Mb resolution and 1,200 regions with 25 kb resolution, are imaged by DNA seqFISH+, followed by sequential IF imaging for 17 chromatin related marks and proteins. Throughout imaging routines, dye conjugated readout probes, which are unique to targets, are hybridized, imaged and stripped off. b, Example image for DNA seqFISH+ in a mESC. Top, DNA seqFISH+ image from one round of hybridization at a single z section. Bottom, DAPI image from the same cell. c, zoomed-in view of the red boxed region in b through five rounds of barcoding in channel 1 decoded with megabase resolution. Images from 16 serial hybridizations are collapsed into a single composite 16-pseudocolor image, which corresponds to one barcoding round. White boxes on pseudocolor spots indicate identified barcodes. d, Zoomed-in view of the yellow boxed region through 60 rounds targeting adjacent regions at 25 kb resolution followed by 20 rounds of chromosome painting in channel 3. Scale bars represent 250 nm in top and bottom zoomed-in images. e, Frequencies of on- and off-target barcodes in channel 1 and 2 per cell. On average, 3,636.0 ± 1,052.6 (median ± standard deviation) on-target barcodes and 14.0 ± 7.4 off-target barcodes are detected per cell (n = 326 cells from the center field of views of the two replicates). f, Average frequencies of individual on-target and off-target barcodes (n = 4,096 barcodes in channel 1 and 2) calculated from n = 326 cells in e, demonstrating the accuracy of the DNA seqFISH+. g, 3D reconstruction of a single mESC nucleus. Left, individual chromosomes labeled in different colors. Middle, reconstruction of two alleles of chromosome 5 colored based on chromosome coordinates. Right, reconstruction of chromosome 8 at 42.7 Mb with 25 kb resolution. h, Concordance between the normalized contact maps from DNA seqFISH+ (upper right) and Hi-C^4^ in mESCs at different length scales. Similar results were obtained for other chromosomes and regions (Extended Data Fig. 3). n = 446 cells from two replicates. i,j. Physical distance as a function of genomic distance Mb resolution (i) and 25 kb resolution (j) H3K27ac enrichments of the entire region are obtained from ChIP-seq^8^. n = 20 chromosome traces in i, j.

DNA seqFISH+ detected 3,636.0 ± 1,052.6 (median ± standard deviation) dots per cell for the barcoded loci targeted at 1 Mb resolution (Fig. 1e Extended Data Fig. 2f), and 5,616.5 ± 1,551.4 (median ± standard deviation) dots per cell in total including 25 kb resolution data (Extended Data Fig. 2g). This corresponds to an estimated detection efficiency of at least 50% in the diploid genome considering the cell cycle phases. The false positive dots, as determined by the barcodes unused in the codebook, were detected at 15.3 ± 7.4 per cell (mean ± standard deviation), about a factor of 200 lower than the actual barcodes (Fig. 1f).

Reconstructed chromosomes (Fig. 1g) in single cells showed clear physical territories for individual chromosomal alleles and have variable structures amongst cells. The DNA seqFISH+ measurements were highly reproducible between biological replicates (Pearson’s R = 0.92 and 0.95 for 1 Mb and 25 kb resolution, respectively) (Extended Data Fig. 2h). We further validated the DNA seqFISH+ data by comparing the pairwise contact maps from the imaging data (Extended Data Fig. 3) to those from the bulk population Hi-C measurement^4^ with a Spearman’s correlation coefficient of 0.89 at 1 Mb resolution and 0.82-0.94 at 25 kb resolution (Fig. 1h and Extended Data Fig. 2i, j), as well as to those from the SPRITE measurement^10^ with a Spearman’s correlation coefficient of 0.83 at 1 Mb resolution (Extended Data Fig. 2k). The genomic versus physical distance scaling relationships for each chromosome are heterogeneous amongst the chromosomes at 1 Mb resolution, as well as at 25 kb resolution for defined regions on each of the chromosomes (Fig. 1i, j). In particular, regions with low H3K27ac marks^8^ tend to have more compact spatial organization (Fig. 1j), suggesting a correlation between physical compaction of the genome and the underlying epigenetic state^29^.

### Integrated measurements of DNA, RNA, and histone modifications in single cells

Mouse ESCs exist in metastable transcriptional states^30–34^. To investigate how this heterogeneity in gene expression relates to chromosome structures and chromatin states within the same single cell, we integrated our analysis of the genome (DNA seqFISH+) with the transcriptome (RNA seqFISH) as well as histone modifications and subnuclear structures (immunofluorescence (IF)) (Fig. 1a and Extended Data Fig. 1a). Briefly, 17 primary antibodies targeting nuclear lamina, nuclear speckle, nucleolus and active and repressive histone modification markers were conjugated with DNA oligonucleotides (oligos) (Fig. 2a and Extended Data Fig. 2d, 5a). These antibodies (see ‘Methods’) and RNA FISH probes for 70 mRNA and introns species were hybridized in the same cells as the DNA seqFISH+ probes. Additionally, 4 repetitive regions that relate to nuclear organization^35,36^ were sequentially imaged, including gene-poor long interspersed nuclear elements (LINE1), gene-rich short interspersed nuclear elements (SINEB1), centromeric minor satellite DNA (MinSat), and telomeres (Extended Data Fig. 5a, b).

**Figure 2.**
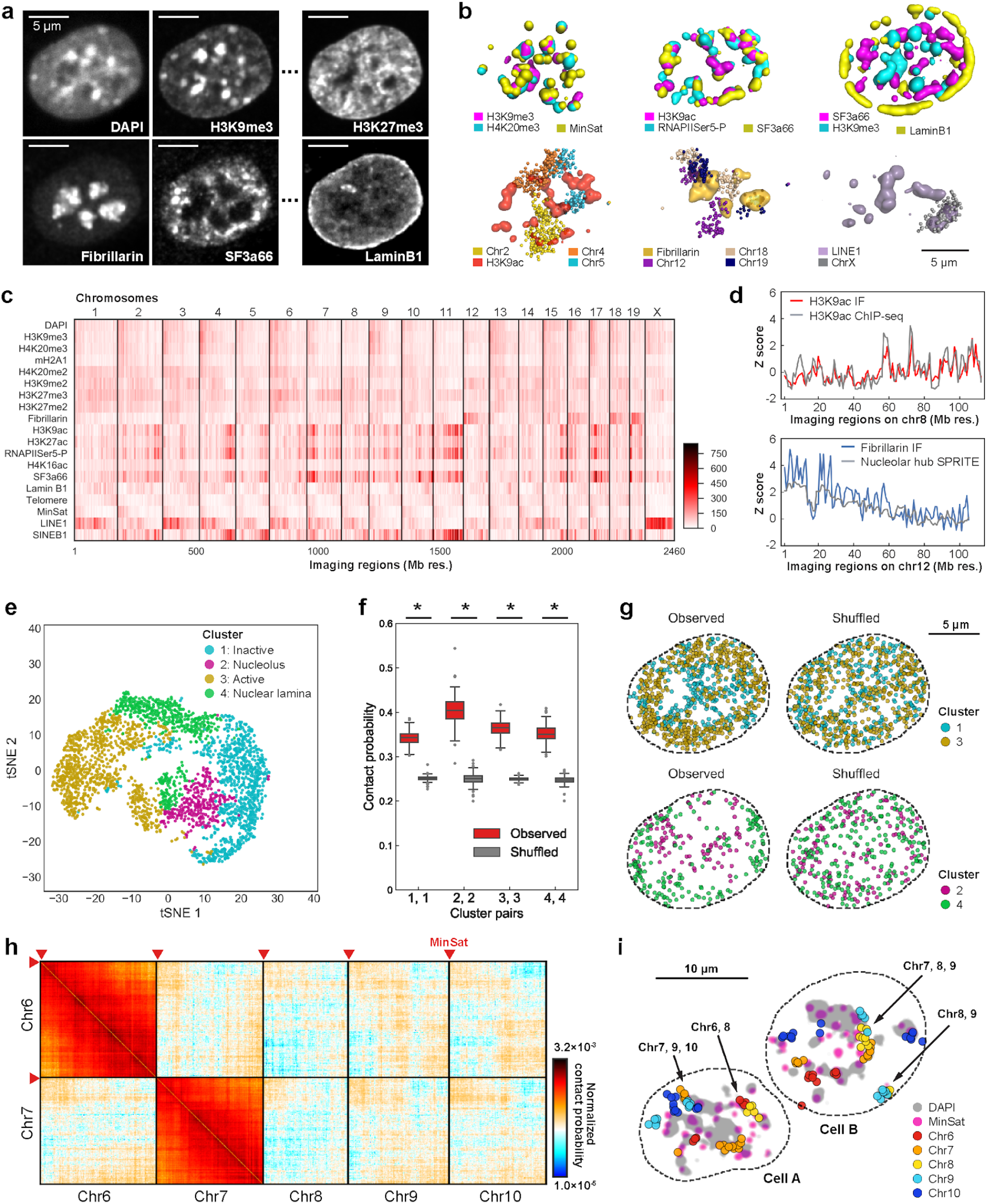
DNA seqFISH+ combined with sequential IF enables multiplexed chromatin profiling in situ. a, images for DAPI and immuno-staining in a mESC nucleus. (n=201 cells) Scale bars: 5 μm. b, 3D reconstructions for sequential immunofluorescence and DNA seqFISH+ in the same cell. Top panels, left, heterochromatin markers (H3K9me3, H4K20me3, MinSat). Middle, transcriptionally active markers (H3K9ac, RNAPIISer5-P, SF3a66). Right, mutually exclusive nuclear stainings (nuclear speckle: SF3a66, heterochromatin: H3K9me3, nuclear lamina: Lamin B1). Bottom panels, left, association between active mark H3K9ac and chr2, 4 and 5. Middle, association between nucleolus marker Fibrillin and chr12, 18 and 19. Right, association between LINE1 repetitive elements and LINE1 rich chrX. c-d, Contact frequencies between DNA seqFISH+ loci (n = 2,460) from 1 Mb resolution data and immunofluorescence markers shown in heatmap (c), comparison with ChIP-seq (H3K9ac)^8^ and SPRITE (nucleolar hub)^10^ results (d). Z scores were computed from all 2,460 loci. e, clustering of the average IF contact profile of individual loci. 4 clusters are shown with inactive (1), nucleolus (2), active (3), and lamin marks enriched (4). n = 2,460 loci (n = 805, 278, 877, 500 loci in each cluster, respectively). f, Probabilities of loci in the same clusters contacting within 1 μm measured in single cells (n = 201 cells) in comparison with those from randomized cluster assignments. *p < 0.001 from a two-sided Wilcoxon’s rank sum test (p = 2.2e-67, 2.3e-67, 2.2e-67, 2.2e-67 from left to right) computed from the distribution of the probabilities in individual cells (n = 201 each). For the boxplots, the center line in the boxes marks median, the upper and lower limits of the boxes mark the interquartile range, the whiskers extend to the farthest data points within 1.5 times the interquartile range, and the gray points mark outliers. g, Mapping of the cluster definitions to DNA seqFISH+ decoded spot location in a single nucleus, for experimental and randomized data. h, Chromosome contact map within 1 μm search radius shows pericentromeric regions near MinSat interact between chromosomes from 201 cells. Colorbar is shown in log-scale. i, Single nuclei reconstruction shows that pericentromeric regions from only a subset of chromosomes interact in DAPI-rich regions in individual nuclei.

We extensively optimized the combined protocols, which included incubating and fixing antibodies with an irreversible crosslinker in addition to the initial formaldehyde fixation, before probe hybridization for RNA seqFISH (see Methods). This optimization allowed us to profile these different modalities without a major loss of signal and accurate alignment between IF and DNA FISH images for over 130 rounds of hybridizations on an automated confocal microscope (Extended Data Fig. 2a-e). For the sequential IF steps, the unique oligos conjugated to individual antibodies were serially read out by hybridizing fluorescently labeled oligos in cycles after DNA seqFISH+ (Extended Data Fig. 2d). RNA species were similarly read out but prior to DNA seqFISH+ (Extended Data Fig. 1a) due to the instability of RNA in heat denaturation. To image target species in all hybridization cycles, we used fluorescently labeled readout oligos (12 to 15-nt), which are then stripped off with 55% formamide to completely remove the signals^27,28^. The multiplexed IF and RNA seqFISH images were then aligned with the DNA seqFISH+ images with 1 voxel (100 nm x 100 nm x 250 nm) resolution.

Nuclear bodies such as the nuclear lamina, nucleoli and nuclear speckles were clearly detected by IF (Fig. 2a, b and Extended Data Fig. 5a). Repressive histone marks (e.g. H3K9me3, H4K20me3) colocalized with DAPI rich regions. and minor satellite DNA (MinSat) corresponded to pericentromeric and centromeric heterochromatin^35,37^ (Fig. 2b top left). Immunofluorescence of RNA polymerase (RNA pol II CTD phospho Ser5) and active marks (H3K9ac) localized to the periphery of nuclear speckles (SF3a66) (Fig. 2b top middle) and were excluded from both heterochromatic regions and the nuclear lamina (Fig. 2b top right). By correlating the IF marks for each pixel, we confirmed that active marks are clustered together and anticorrelate with repressive marks (Extended Data Fig. 5c).

To validate the integrated multiplexed IF and DNA seqFISH+ data, we generated a chromatin profile for each DNA locus by calculating the contact frequencies with individual IF markers as well as repetitive elements (Fig. 2c). We compared this profile with ChIP-seq data^8^ and found strong agreements (Fig. 2d top) with Pearson’s R = 0.77 (H3K9ac). In addition, highly expressed loci, such as genes encoding tubulin and ribosomal protein subunits, were within regions enriched for active marks and nuclear speckle (Supplementary Table 3), agreeing with the active hubs found by SPRITE^10^ (Fig. 2d bottom). Chromosomes 2, 4 and 5, which appear in the active hubs in the SPRITE data, were spatially close to H3K9ac-enriched regions in single cells (Fig. 2b bottom left). Chromosomes 12, 18 and 19, which contain rDNA units, showed significant association with the nucleoli (Fig. 2b bottom middle, 2c), consistent with the nucleolar hub assignments by SPRITE^10^. In addition, DNA seqFISH+ loci with high LINE1 FISH intensity pixels localized mainly to the LINE1-rich X chromosome (Fig. 2b bottom right, 2c)^38^.

To determine whether loci with similar chromatin profiles aggregated in single cells, we first grouped the loci into four clusters based on the IF data (Fig. 2e). By comparing the chromatin marks enriched in individual clusters (Extended Data Fig. 6c), we annotated cluster 1 as “inactive”, cluster 2 as “nucleolus”, cluster 3 as “active”, and cluster 4 as “nuclear lamina”. We then mapped the clusters within individual cells and measured to what extent clusters intermingle. Briefly, we determined how frequently loci with the same or different annotation were within 1 μm of each other (defined as the ‘contact probability’, Fig. 2f and Extended Data Fig. 6a). We found that loci within the same cluster display a higher contact probability than predicted by a random distribution (Fig. 2f). However, we also observed significant intermingling of loci from different clusters in single cells (Fig. 2g and Extended Data Fig. 6b). In addition, we obtained A/B compartment assignments of chromatin from bulk population Hi-C measurements^1,4^ and found that A/B compartments also intermingle in single cells (Extended Data Fig. 6d-g).

The contact maps generated from DNA seqFISH+ revealed that pericentromeric regions between different chromosomes interact (Fig. 2h and Extended Data Fig. 6h), consistent with the observation from Hi-C^39,40^. In single cells, we found that regions near the starting coordinates of individual chromosomes (at 3-8 Mb in each chromosome) are physically close to MinSat-rich regions and DAPI-rich regions of the nucleus (Fig. 2i). Importantly, our data showed that these interactions are variable in single cells: the non-repetitive regions (at 3-8 Mb in each chromosome) from different subsets of chromosomes aggregate near DAPI-rich pericentromeric and centromeric heterochromatin (Fig. 2i, compare ‘cell A’ with ‘cell B’, Extended Data Fig. 6i). When these data were averaged over all cells in the ensemble contact map, they showed that regions near the starting coordinates on every chromosome contact every other pericentromeric region (Fig. 2h, Extended Data Figure 6h). Taken together, the bulk measurements explained global trends of interchromosomal interactions, but the chromosome organizations in single cells were heterogeneous and did not completely recapitulate global trends.

### Combinatorial IF marks define nuclear “zones”

To analyze the IF data fully at the subcellular level in single cells^41^, we clustered individual binned voxels (200 nm x 200 nm x 250 nm) within the nuclei, based on their combinatorial chromatin profiles (Fig. 3a). There were at least 12 major clusters with distinct combinatorial IF patterns, as represented by Uniform manifold approximation and projection (UMAP)^42^ (Fig. 3b). 15 chromatin marks, excluding cell cycle makers, and DAPI were used in this analysis. Some of these distinct clusters, or nuclear “zones”, corresponded to known nuclear bodies such as the nuclear speckles (cluster 1), enriched with the splicing factor SF3a66; the nucleolus (clusters 8 and 9) enriched with Fibrillarin, a key nucleolar protein, and nuclear lamina (cluster 10) enriched with Lamin B1 (Fig. 3c and Extended Data Fig. 7a). In addition, functional zones (cluster 2) enriched in active marks (RNAPII ser5 phosphorylation and histone acetylation marks) form contiguous regions in the nucleus that often surround the nuclear speckles. The three heterochromatin zones (clusters 5, 6 and 7) had distinct combinatorial chromatin marks (Fig. 3c). In addition, several zones showed a mixture of marks, such as cluster 3 and 4 with mixed repressive and active marks, and cluster 12 with mixed active and nuclear lamina marks (Fig. 3c). By mapping these zones back to the images, we confirmed that they form physically distinct regions in single nuclei (Fig. 3d and Extended Data Fig. 7b, c), rather than well mixed in the nucleus, suggesting that zones may form due to phase separation^43–45^ of different nuclear components.

**Figure 3.**
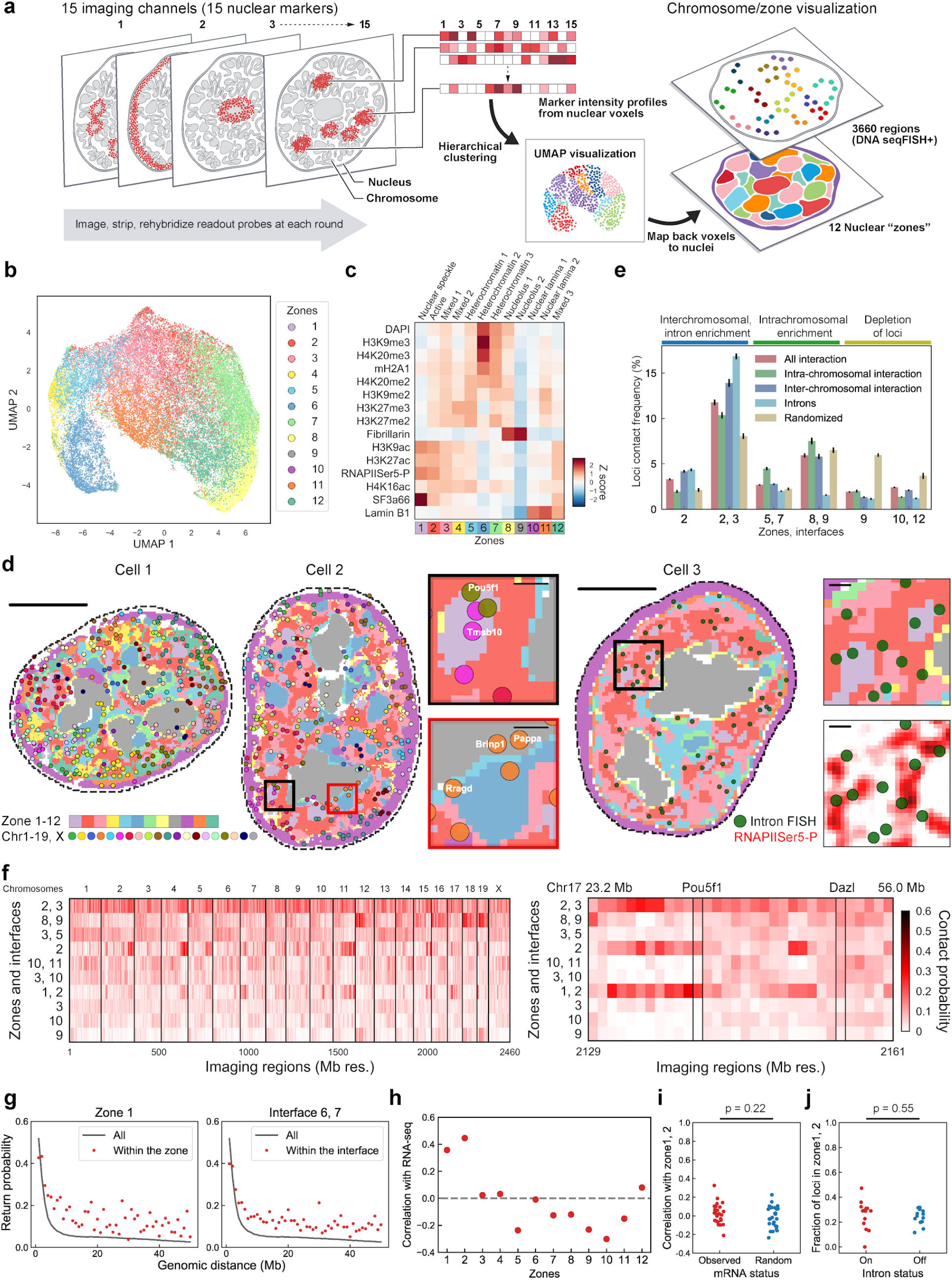
Combinatorial subcellular chromatin patterns reveal distinct nuclear zones and their differential association to DNA loci in single cells. a, Analysis workflow for the pixel-based combinatorial chromatin profiling. Individual pixels with the 15 chromatin markers imaged are clustered with hierarchical clustering and visually represented by UMAP. Voxels from individual clusters or “zones” are mapped back to individual nuclei, and overlaid with DNA seqFISH+ dots. b, UMAP representation for 44,000 pixels sampled from 201 cells, labeled with 12 zones. c, Heatmap for differential enrichment of individual chromatin markers in each zone computed from 201 cells. d, Cells 1 and 2, reconstruction for 12 nuclear zones and DNA loci from 20 chromosomes at a single z plane. Zoomed-in view of the black and red boxed regions represent inter-chromosomal localization of gene loci such as Pou5f1 in active zone 1, and intra-chromosomal localization around heterochromatin zone 6 and nucleolar zone 9, respectively. Cell 3, nuclear zones with 1,000 gene intron FISH. top right panel, zoomed-in view of black boxed region, showing the enrichment of nascent transcription active sites at the interfaces of nuclear zones. bottom right panel, same view with nascent transcription active sites are enriched at the edge of the RNAPIISer5-P staining (background-subtracted), colored by intensity.(n = 172 cells; Scale bars: 5 μm). Additional cells are shown in Extended Data Fig. 7c, d. e, Contact frequency between DNA loci and zones/interfaces in single cells, calculated for all loci, loci contacting other loci from the same chromosomes (intra-chromosomal), interchromosomal, transcriptional active sites measured by intron FISH and random loci. Error bars are standard errors over 20 bootstrap trials. f, Heatmap for contact probability between DNA loci, nuclear zones and interfaces for 2,460 loci in 1 Mb resolution data (n = 201 cells). g, Return probability as a function of genomic distance, showing higher return probability for loci within a zone or interface. h, Pearson correlation of zone enrichment and RNA expression level for all 1 Mb resolution loci (n = 2,460 loci). i, Correlation between mRNA counts of the profiled genes and their association to active zones (1, 2) in single cells in comparison with randomized samples. Each dot represents a gene (27 genes, n = 125 cells). The genes with mean copy numbers of > 2 per cell were used. Cells in the center field of views were used. j, Comparison between intron state and active zone (1, 2) association of the corresponding alleles (14 genes, n = 201 cells) The genes with mean burst frequencies of > 0.1 per cell were used. P values from a two-sided Wilcoxon’s rank sum test between different mRNA or intron status of the genes are shown in i, j.

### Active loci are enriched in the interfaces between zones

We overlaid the DNA seqFISH+ loci with the nuclear zone assignments, and found that DNA loci were enriched at the interfaces between zones (Fig. 3d, e and Extended Data Fig. 7c, e, Supplementary Table 4). For instance, the probability of detecting DNA loci at the interface between active (zone 2) and mixed zones (zone 3) was 46.3% higher than by random chance (Fig. 3e). In particular, interchromosomal interactions were enriched at the zone 2/3 interface and within zone 2 (Fig. 3e), whereas intrachromosomal interactions were enriched at interfaces of heterochromatin zones (interface of zone 5/7) and nucleolus zones (interface of zones 8/9). In contrast, DNA loci are depleted from the nucleolus zone (zone 9) and nuclear lamina interfaces (interface of zones 10/12) (Fig. 3e), suggesting different physical properties of the nuclear zones and interfaces in chromosome organization.

To detect the nascent transcription active sites on the chromosomes, we imaged intronic RNAs from 1,000 genes^27^ and performed multiplexed IF in the same cells. We found that transcriptionally active sites are close to the nuclear speckle zone (Fig. 3e), which is consistent with speckles being enriched for splicing machinery. Transcriptional active sites were also enriched in the interface between active and mixed zones (Fig. 3e and Extended Data Fig. 7e); their contact frequency at this interface was two-fold higher than predicted by random chance. Intriguingly, active loci were also observed at heterochromatic and nucleolus zone interfaces with lower frequency (Fig. 3e and Extended Data Fig. 7e). Furthermore, active loci appear at the edges, rather than the center, of RNAPII dense regions in the nuclei (Fig. 3d and Extended Data Fig. 7d). These results agree with our previous finding that transcriptionally active loci are dispersed in the nucleus and have significant interchromosomal interactions^27^. Overall, our DNA seqFISH+ and intron data with the combinatorially defined IF zones indicate that a significant fraction of each chromosome localizes to a 2D shell in the interface regions in between nuclear zones, whereas satellite repeat regions, telomeres and rDNA repeats are packed into or around nuclear bodies such as heterochromatin and nucleolus, respectively.

### Specific loci have stereotypical but heterogeneous zone association in single cells

To examine the relationship between the physical locations of specific DNA loci and their proximity to specific zones or interfaces, we computed their contact probabilities and found that each locus had stereotypical zone associations, but were also highly variable among cells (Fig. 3f, Extended Data Fig. 7f). For example, the Pou5f1 (Oct4) locus was predominantly associated with active zones and interfaces (zones 1, 2, 3), but at a lower frequency was near the nucleolus zones and interfaces (zones 8, 9) (Fig 3f, right). In contrast, the Dazl locus was associated with active, heterochromatic and nuclear lamina zones and interfaces with similar frequencies (Fig. 3f right).

Neighboring loci within 1 Mb tended to be associated with similar nuclear zones (Extended Data Fig. 8a), suggesting that megabase regions of the chromosomes are stably associated with particular nuclear bodies or zones. To determine whether loci with similar zone assignments are more likely to be spatially close regardless of genomic distance, we examined the likelihood that pairs of loci are found within a 500 nm radius sphere in single cells as a function of genomic distance (which we term “return probability”). The return probability for all loci along the entire chromosome decreased rapidly, and loci separated by >10 Mb had only a 6.0% probability of remaining within the sphere. However, loci with the same zone assignments had a return probability as high as 13.4% (zone 1) and 17.9% (zone 6/7 interface) even at 20 Mb distance (Fig. 3g). Thus, at the single cell level, even though loci within a few Mbs tend to move far away, loci sometimes loop back to the same nuclear zone even across long genomic distances (Extended Data Fig. 8b, e).

### Zone associations correlate with gene expression in bulk but not at the single cell level

We next asked whether the proximity to specific zones are functionally relevant and predictive of expression. For each locus, we correlated its frequency of association with a zone to its log-expression level (Fig. 3h). An association with active and nuclear speckle zones, and the interface regions between active, speckle and mixed zones, showed a high correlation with expression, in both 1 Mb (Fig. 3h and Extended Data Fig. 8c). Conversely, an association with lamin, heterochromatic and nucleolar zones negatively correlated with expression (Fig. 3h). These results based on imaging data are consistent with previous findings from DamID and TSA-seq^7,9^. Active zones also correlated with the density of RNA polymerases on the loci, as measured by GRO-seq^46^ and to early replication domains measured by Repli-seq^47^ (Extended Data Fig. 8c). To measure how well zone proximity predicts gene expression, we constructed a linear model (supplementary material) and found that log-expression levels are predicted with Pearson’s R = 0.57 (Extended Data Fig. 8d). We note that although mean expression is anticorrelated with nuclear lamina and nucleolar association, we still observed some active loci and active transcription sites (Fig. 3d, f) around the nuclear lamina and nucleolus^27^, indicating that this correlation has exceptions at the single cell level.

Unexpectedly, we observed no correlation between mRNA and intron expression and proximity with active and speckle zones in single cells amongst the genes we examined (27 genes for mRNA spanning a large range of expression levels; 14 genes for intron) (Fig. 3i, j). In other words, a highly expressed gene tends to be close to active/speckle zones in all cells regardless of its expression and bursting state in individual cells. Similarly, a gene that is on average lowly expressed tends to associate with heterochromatin and lamin zone even in the subset of cells where it is bursting. Given the lifetime of mRNA and introns, which are on the order of several hours and minutes respectively, it is possible that chromosomal positions are relatively stable for a given cell state, leading to an apparent lack of correlation with the stochastic bursts of RNAs production in snapshot measurements of single cells.

### Global chromatin states are heterogeneous in mESCs

The overall intensities of IF signals in the nucleus showed substantial heterogeneity among single cells (Fig. 4a). Clustering analysis of the IF data shows at least 7 distinct states based on global chromatin modification levels, as viewed by heatmap (Fig. 4b) and UMAP representation (Fig. 4c). These states identify previously unexplored combinatorial patterns of global chromatin or protein marker levels in single cells, as well as co-regulation of those markers (Extended Data Fig. 9c). To examine the relationship between these IF states and the transcriptional states at the single cell level, we used previously characterized markers of mESC phenotypes (Supplementary Table 5). Specifically, we analyzed naive or pluripotency markers (Zfp42, Esrrb, Nanog and Tbx3) and primed markers (Otx2, Dnmt3b and Lin28a), which clustered as expected (Extended Data Fig. 9a, b). Interestingly, IF states only partially overlapped with transcriptional states. For example, Zfp42, Nanog and Esrrb expressing cells in the “ground” pluripotent state, as well as Otx2 expressing cells in the orthogonal “primed” states are present in most IF clusters (Fig. 4b). Conversely, clusters based on RNA states (Extended Data Fig. 9a, b) contain a mixture of cells with distinct chromatin states. These results suggest that chromatin states, defined by combinatorial profiles of the overall abundance of IF marks in single cells, adds to the complexity in the mESCs states, beyond the previously characterized heterogeneity in transcriptional states^30–34^ and are independent from cell-cycle phases (Extended Data Fig. 9d, e).

**Figure 4.**
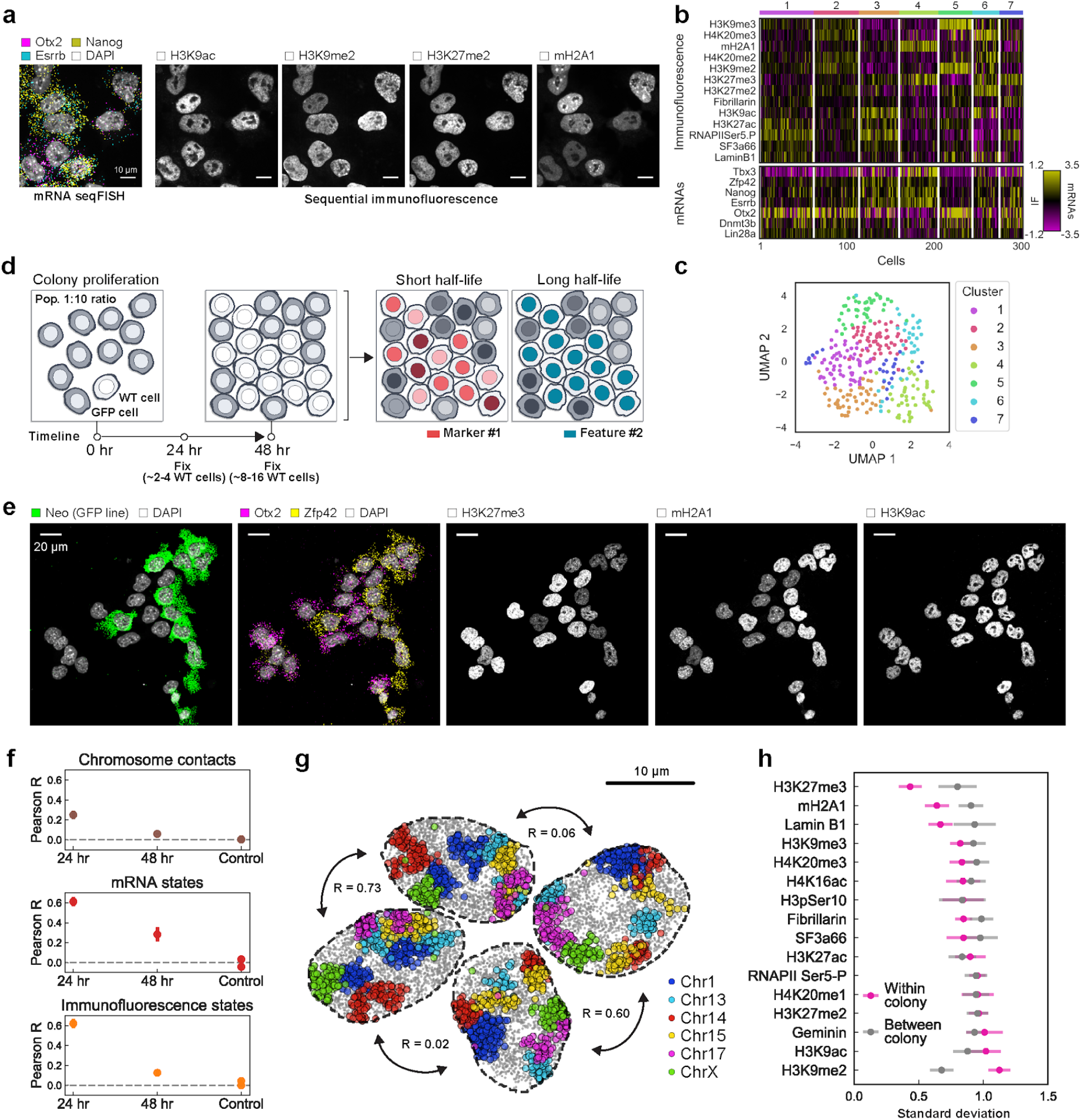
Global chromatin states are highly variable and dynamic in single cells. a, The intensities of multiple IF markers show large heterogeneities in single cells. mRNAs for Essrb, Nanog and Otx2 are shown in the same cells. Images are from the same single z section. Scale bars: 10 μm. b, Heatmap of cell clusters with distinct IF profiles. bimodally expressed Nanog, Zfp42, Tbx3 are distributed over several IF clusters. c, UMAP representations of the IF clusters. n = 302 cells in the center field of views from two replicates in b, c. d, Schematic of single colony tracing experiments. Unlabeled mESCs and GFP/neomycin-resistance gene (Neo) expressing mESCs are mixed with 1:10 ratio and seeded on coverslips. Cells were fixed at different time points, where unlabeled cells are expected to form 2-4 cell colonies at 24 hours and 8-16 cell colonies at 48 hours. Intensity of markers with fast dynamics are expected to be heterogeneous within a colony while those of long memories would be more homogeneous. e, Representative maximum intensity z-projected images of sequential RNA FISH and IF for one 48-hour colony, showing expression level heterogeneities for RNA and chromatin markers within and between the colony of unlabeled cells and the surrounding unrelated GFP cells. Scale bars: 20 μm. f, Mean Pearson correlation between cells within colonies decays slowly for mRNA and chromatin states, and quickly for chromosome conformations. Control measures correlation between colonies for both 24- and 48-hour datasets. Error bars are standard error over 20 bootstrap trials. n = 53, 117 unlabeled cells within colonies in 24-hour and 48-hour dataset. g, Chromosomes reconstructions for unlabeled cells from a 24-hour colony shows similarities between two sets of neighboring cells (maximum z projection). 6 chromosomes are shown for visual clarity. h, Standard deviation of individual IF marker intensities in 48-hour colonies compared to those between colonies. H3K27me3, mH2A1 and Lamin B1 have less variance in cells within a colony, which can be seen in e. Error bars are standard errors over 20 bootstrap trials. n = 117 unlabeled cells within colonies.

### Chromatin states can be inherited over multiple generations

To examine whether the heterogeneity in chromatin states, expression and chromosome organization are stable or are dynamic over generations, we performed clonal analysis experiments. Because DNA/RNA seqFISH and multiplexed IF cannot be performed in live cells, clonal analysis allows us to infer the dynamics of chromatin and RNA states^34^: if clonally related cells have similar molecular states, then those states are likely to have slow dynamics, and vice versa (Fig. 4d). We seeded unlabeled mESCs among GFP-positive mESCs at a 1:10 ratio and cultured them for 24 and 48 hours, which are approximately 2 and 4 generations respectively, such that each unlabeled mESC colony likely arises from a single cell (Fig. 4d, e and Extended Data Fig. 10a). We correlated mRNA states, IF states and chromosome contacts within each unlabeled colony fixed at either 24 and 48 hours (Fig. 4f) and excluded the GFP-positive cells from the analysis.

Interestingly, we found that chromosome contacts correlated among cells within a colony but not among those from randomized cell assignments (Fig. 4f and Extended Data Fig. 10b, c). Moreover, chromosome contacts in single cells were highly correlated between pairs of cells that were physically close, possibly sister cells, and are mostly uncorrelated with other cells in the colonies (Fig. 4g). These data suggest that chromosome contacts are preserved across one cell cycle between sisters, which supports previous observations that the global arrangement of chromosomes is heritable for one generation^48^, but are then rapidly lost after 2 generations.

In contrast, overall mRNA and chromatin profiles were highly correlated amongst most cells within a colony (Fig. 4f), and maintained some correlation even at the 48hr time points. As an internal validation, we examined the expression of pluripotency genes such as Nanog and Tbx3 and found that they were highly correlated within colonies (Extended Data Fig. 10d), consistent with previous reports that their expression states propagated for multiple generations in mESCs^31,34^.

Finally, we examined the dynamics of individual IF markers rather than global chromatin states. Specifically, we calculated the variance of IF intensities within each colony compared to between colonies (Fig. 4h). Notably, some IF marks, such as mH2A1 and H3K27me3, were highly correlated within colonies but not between colonies, suggesting that these chromatin features are heritable across at least 3-4 generations (Fig. 4e). On the other hand, many IF marks, such as H3K9ac, did not correlate within a colony nor between colonies, suggesting that these features are rapidly fluctuating. Single pluripotent stem cells exhibit highly variable responses to differentiation signals^49–51^, with some cells highly resistant to differentiation even when lineage specific transcription factors are expressed. The heterogeneous long-lived chromatin features could represent a “hidden variable” in differentiation experiments^49–52^, warranting furthering investigation with imaging-based lineage tracing tools such as MEMOIR^53^.

## Discussion

Our integrated spatial genomics approach with DNA seqFISH+ along with multiplexed IF and RNA seqFISH enables multimodal profiling of chromosome structures, chromatin states, and gene expression within the same single cells. Our discovery that DNA loci, especially active gene loci, reside mostly at zone interfaces relied on the visualization of imaging data that span multiple modalities and covers genomic scales. These findings raise further questions about the nature of the nuclear zones. Biophysically, are they liquid droplets of proteins that phase separate^43^? Functionally, do regulatory factors diffuse in 2D or 3D to search for their target genes, and are their kinetics accelerated in these interfaces compared to in bulk^54^? Furthermore, given the presence of the nuclear zone and the large variability even in the total abundance of individual IF marks in single cells, it is striking to notice the concordance between the population averaged ChIP-seq data, which measures molecular interactions, and the “chromatin profiles” (Fig. 2c) generated by averaging single cells in the imaging-based data, which measures physical proximity between large nuclear zones and DNA loci. These interactions between DNA loci and nuclear zones/interfaces might represent a new layer of regulation, beyond specific binding sequences. Ultimately, the observation of heterogeneous and long-lived global chromatin states raises the question of whether these states have distinct pluripotency and differentiation potentials and reveals additional complexity in cell states beyond those previously appreciated from transcriptome profiling. We anticipate that the integrated spatial genomics approaches will enable further exploration of those questions in many biological contexts.

## Author contributions

Y.T. and L.C. conceived the idea and designed experiments. Y.T. designed probes with help from J.T. and C.-H.L.E. Y.T. and J.Y. prepared and validated all the experimental materials. Y.T. performed all the experiments with help from J.Y. Y.T. and N.P. performed image analysis with help from J.W. and S.S. Y.T. and L.C. analyzed data with N.O., S.Z. L.C., M.G. and G.-C.Y. supervised the analysis process. Y.T. and L.C. wrote the manuscript with input from C.-H.L.E. and G.-C.Y. L.C. supervised all aspects of the projects.

## Acknowledgment

We thank I. Strazhnik for help with the figures and A. Anderson for help with the manuscript. C. Karp for custom made flowcells and H.-J. Ahn for the early phase of the antibody conjugation. B. Bonev for the Hi-C data. This project is funded by NIH 4DN DA047732 and supplement, and Paul G. Allen Frontiers Foundation Discovery Center.

## Competing Interest

L.C. is a co-founder of Spatial Genomics Inc.

---

**Extended Data Fig. 1.**
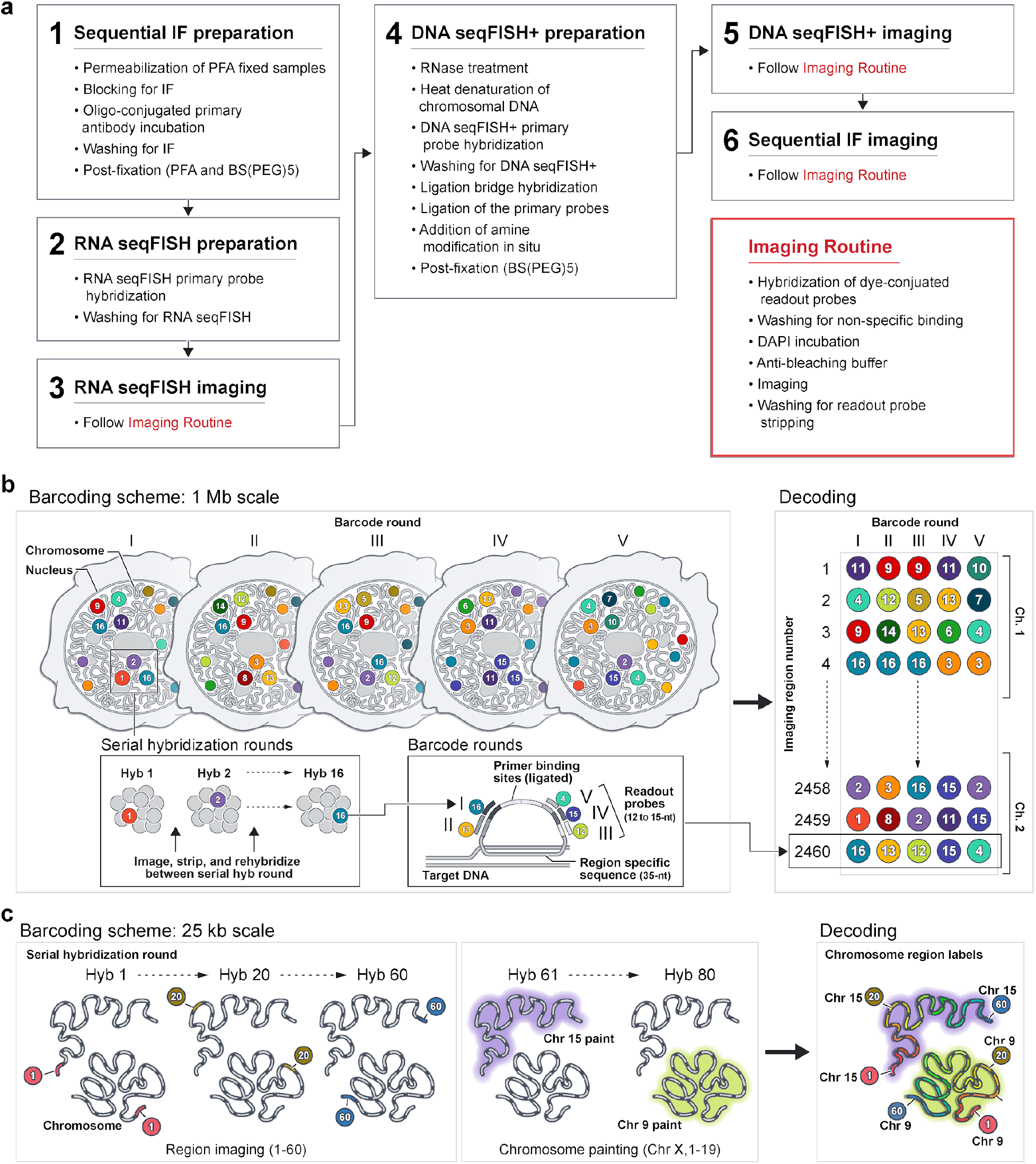
Detailed schematics of the integrated spatial genomics analysis with DNA seqFISH+, RNA and intron seqFISH and multiplexed immunofluorescence. a, Flow chart of the experimental procedures. Samples are fixed with PFA, followed by oligo-conjugated primary antibody hybridization, post-fixation with PFA and BS(PEG)5, and RNA seqFISH, then samples are hybridized with DNA FISH. This optimized protocol ensures good alignment between DNA seqFISH+ data with the multiplexed IF data on a voxel by voxel level. b, Schematics of DNA seqFISH+ for the 1 Mb resolution dataset. 5 round of barcoding allows over 2,048 barcodes to be detected with 2 rounds of dropout error correction in each fluorescent channel. Two fluorescent channels are used to cover a total of 2,460 loci, spaced approximately 1Mb apart in the genome. In each round of barcoding, 16 rounds of hybridization are performed to generate 16 pseudocolors. DNA dots detected in each pseudocolor channel are fitted in 3D to determine their centroid location with super-resolution and compiled across all 16 pseudocolors to generate a super-resolved image. Over 5 rounds of barcoding (overall 80 rounds of serial hybridizations), the identity of all DNA loci are decoded. Every DNA loci should appear once in every barcoding round in a single pseudocolor. The barcoding table is shown on the right. DNA FISH probes contain all 5 rounds of barcode readout sequences. Each sequence has a possible choice of 16 sequences, corresponding to one of the pseudocolors. For each gene, 5 out of the 80 hybridizations will result in hybridization events and fluorescent readout probes bound on the primary DNA hybridizing probes. To preserve the DNA primary probe on the chromosome over all 80 rounds of hybridizations, the primary probes are padlocked onto the chromosomes by T4 DNA ligase after the initial hybridization. c, Barcode scheme for the 25 kb resolution seqFISH+ data. 60 adjacent 25 kb regions are sequentially readout and imaged in 60 rounds of hybridization. This is carried out in parallel on 20 chromosomes. In other words, each round of hybridization images 20 different loci on different chromosomes. An additional 20 rounds of hybridization were carried out to label each chromosome one at a time to assign chromosomal identity to each loci individually. The 1 Mb resolution data were collected in the 640-nm (channel 1) and 561-nm (channel 2) channels, while the 25 kb resolution data was collected in the 488-nm channel (channel 3).

**Extended Data Fig. 2.**
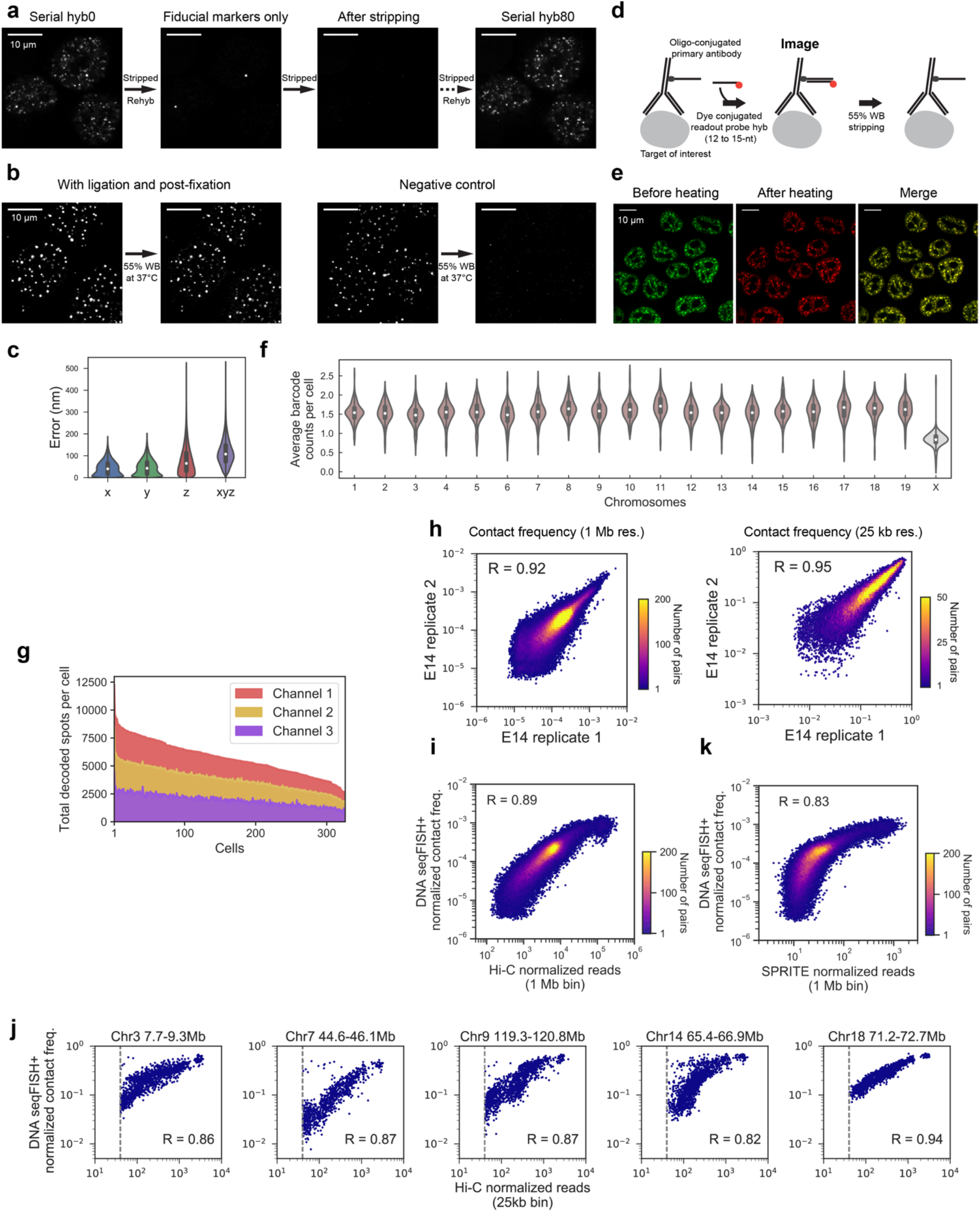
Optimization and validation for DNA seqFISH+. a, Primary probes are still bound after more than 81 rounds of hybridization and the specific signals return. Fiducial markers targeting a repetitive region of the genome were also imaged initially for an alignment. b, Ligation of primary probes prevents dissociation during readout probe stripping step. 55% formamide wash buffer (WB) solution at 37°C was added to the cells for 16 hours with and without the primary probes padlocked to the DNA. Probes are retained in the ligated sample, and not retained in the unligated sample. Note that 55% WB are used at room temperature for 2 minutes at each time during seqFISH routines, which is a less stringent condition compared to the validation here. c, localization errors of fiducial markers across hyb 1 to 80, n = 71,981 aligned spots for x, y and n = 87,879 aligned spots for z. For x and y alignments, we filtered out aligned dots that were more than 2 standard deviations away from the mean displacement at each hybridization, and new alignments are computed. d, Schematics of the multiplexed IF experiments with oligo-conjugated primary antibodies. Readout probes are used to hybridize, image, and strip with 55% formamide to target different antibodies with a unique readout sequence in each round of hybridization. e, Preservation of the nuclear structure through the double fixation procedure. Good colocalization (yellow in the right panel) of the nuclear speckles (SF3a66) before and after heating. f, The total number of dots detected for each loci per cell across all 20 chromosomes. Note that 2 dots per cell are not 100% detection efficiency because some cells are in G2 (4 alleles in total). X chromosome has half the number of dots detected per loci (0.84 ± 0.21 (median ± standard deviation)) compared to the other autosomes (1.57 ± 0.27), because E14 mESC is a male diploid cell line. g, The total number of dots detected in each of the fluorescent channels in single cells. Channels 1 and 2 contain the 1 Mb loci and channel 3 contains the 25 kb data. n = 326 cells from the center field of views of the two replicates in f, g. h, Comparisons of the two replicates of DNA seqFISH+ experiments’ contact frequencies for all unique intra-chromosomal pairs of loci for the 1 Mb (n = 2,460 loci) and 25 kb data (n = 1,200 loci). n = 201, 245 cells in the replicates. i, Comparison of intra-chromosomal contact frequencies for the 1 Mb loci in autosomes between DNA seqFISH+ and Hi-C^4^. Spearman’s R = 0.89 computed from n = 84,707 unique intra-chromosomal pairwise combinations. Hi-C data were binned with 1 Mb, and overlapping regions within 1 Mb were excluded from this analysis. j, Comparison between DNA seqFISH+ and Hi-C for the 25 kb loci. Median Hi-C reads vary depending on the 1.5 Mb chromosomal regions targeted, ranging from 0.9 to 203.2. We used 5 chromosomal regions with Hi-C reads greater than 40 per 25 kb bin for comparison, showing Spearman’s R ranged from 0.82 to 0.94 computed from n = 948-1,776 unique pairwise combinations. k, Comparison of contact frequencies for the 1 Mb loci in autosomes between DNA seqFISH+ and SPRITE^10^, Spearman’s R = 0.83. The same binning and filtering were used as the Hi-C analysis in i. n = 446 cells from two replicates in i-k.

**Extended Data Fig. 3.**
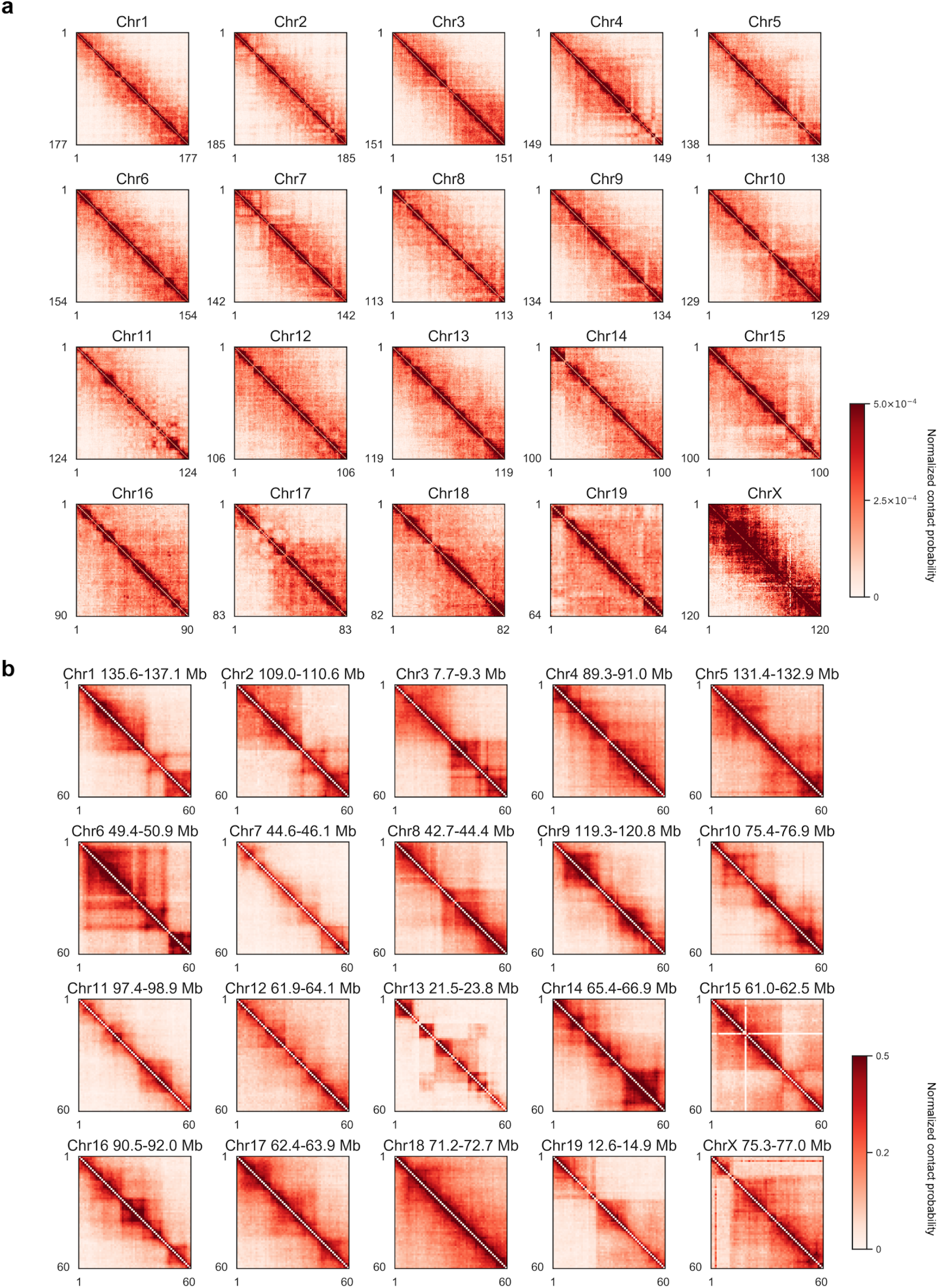
Pairwise chromosome contact maps by DNA seqFISH+. a, For chromosomes 1-20 in the 1 Mb resolution data. b, For the 20 regions measured in the 25 kb data. X- and Y-axis represent the number of loci imaged in each chromosome, and n = 446 cells in a, b.

**Extended Data Fig. 4.**
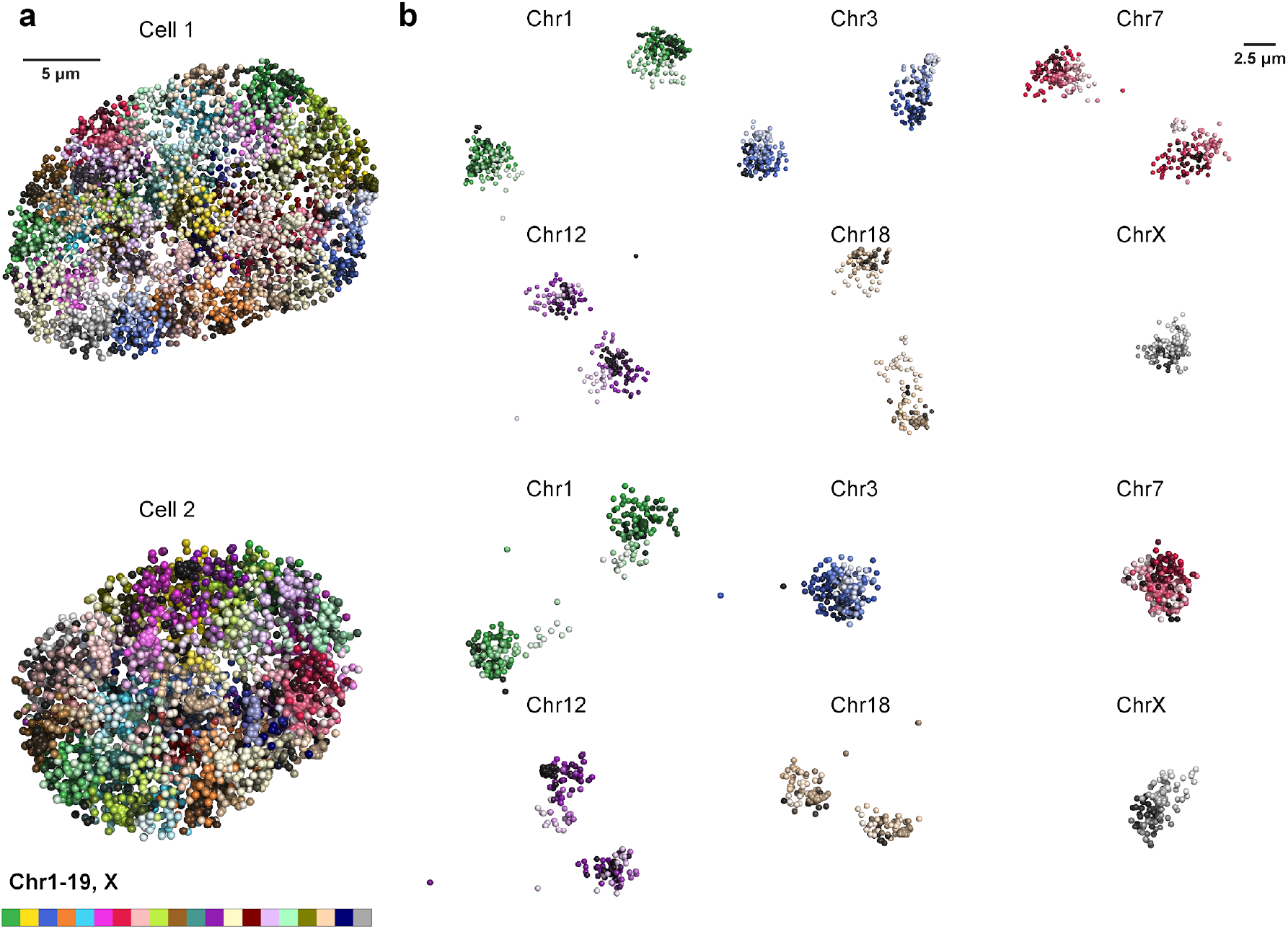
Visualization for DNA seqFISH+ in single nuclei. a, 3D reconstruction of a single mESC nuclei with individual chromosomes labeled in different colors. Cell 1 is shown in Fig. 1 and Cell 2 is shown in Fig. 2. b, 3D reconstruction of individual chromosomes, colored based on chromosome coordinates (light to dark colors). Chromosomes are from cell 1 and cell 2 in a.

**Extended Data Fig. 5.**
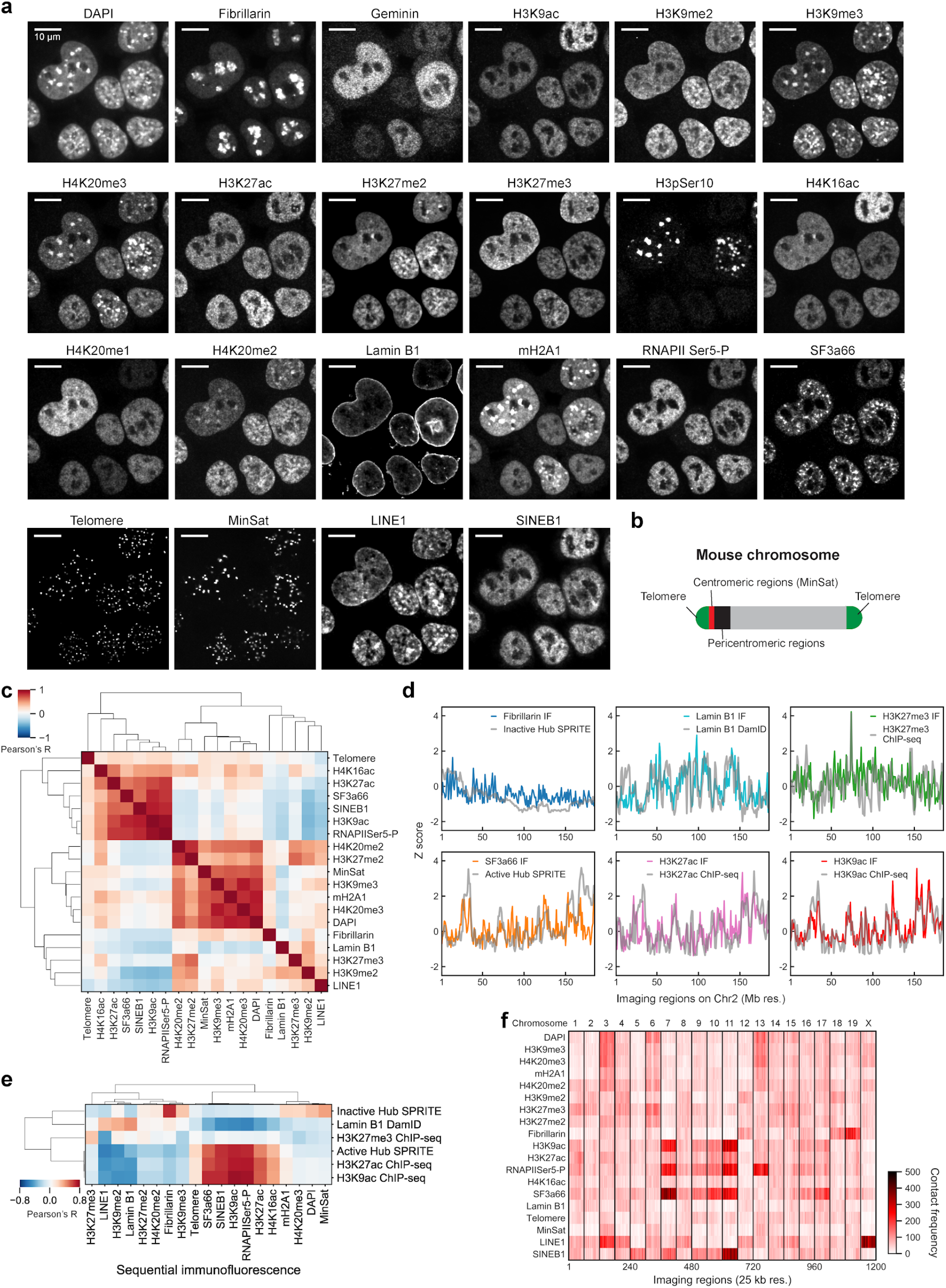
Validation for sequential immunofluorescence and repetitive element imaging. a, 17 antibodies and 4 repetitive elements are imaged along with DAPI. Individual cells have different patterns of IF staining. Note the DAPI patterns are not identical between cells. Similarly, marks that are colocalized with DAPI-rich pericentromeric heterochromatin regions are different between cells and even between different pericentromeric regions in a single cell. b, Schematic showing the position of repetitive regions (telomere and MinSat) along the chromosome. c, Correlation matrix of the different IF markers based on their association with different loci (Fig 2c), computed from 2,460 loci with 1 Mb resolution data (n = 201 cells). d, additional examples for contact frequencies between DNA seqFISH+ loci from 1 Mb resolution data and immunofluorescence markers in comparison with ChIP-seq^8^, SPRITE^10^ and DamID^7^. Z scores were computed from all 2,460 loci. e, Correlation between the DNA seqFISH+ and IF data with other methods^7,8,10^ in measuring chromatin binding. 1 Mb DNA seqFISH+ data are used and the reference data are binned with 1 Mb. Pearson’s R values were computed from the averaged enrichments of individual markers on the 2,460 loci (n = 201 cells). f, Contact frequencies between DNA seqFISH+ data (n = 1,200 loci) with 25 kb resolution and immunofluorescence markers shown in heatmap (n = 201 cells).

**Extended Data Fig. 6.**
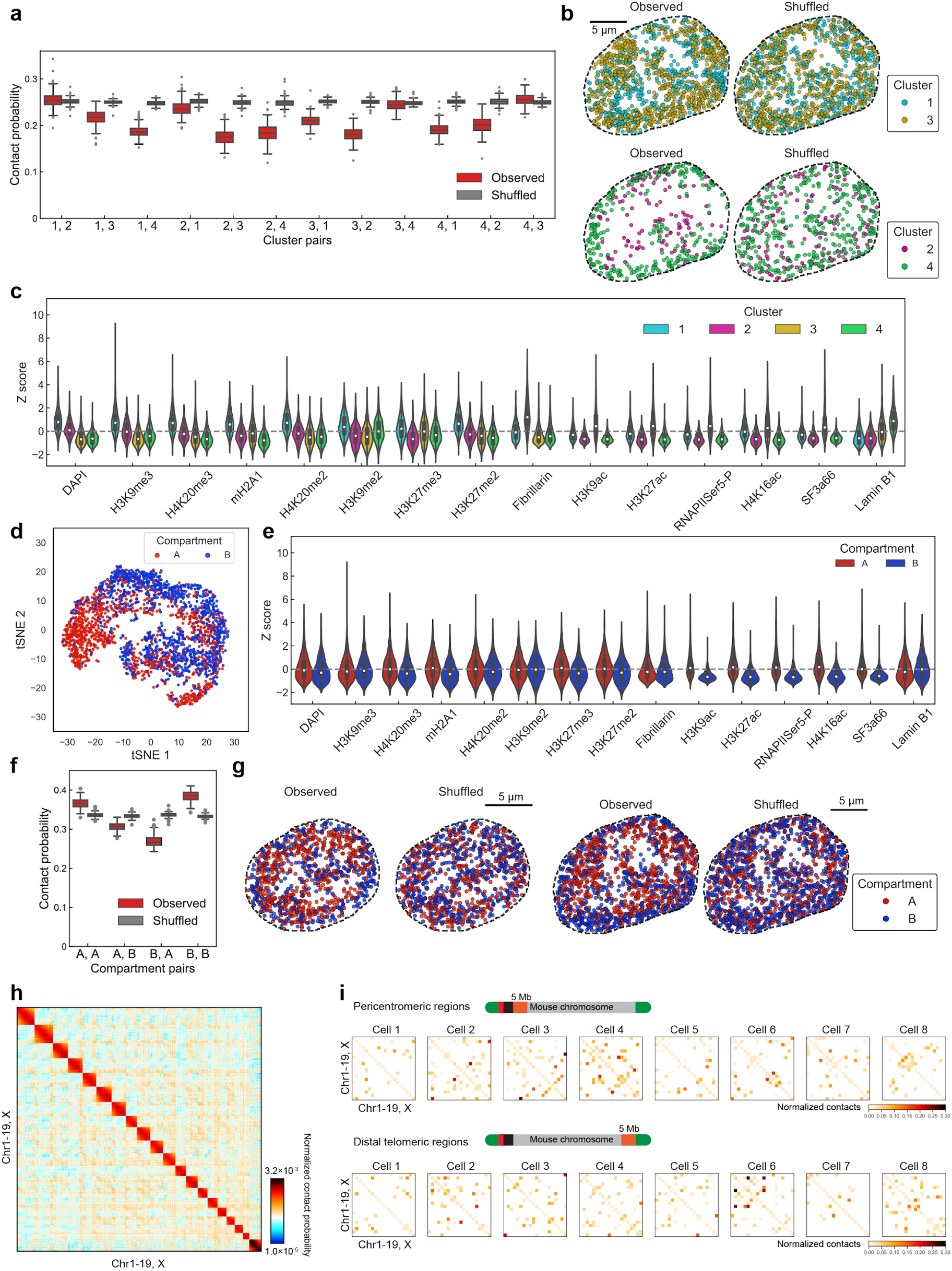
Additional comparison between population level and single cell level chromosome organization. a, The contact probability between loci in one cluster with other clusters in single cells. Clusters are defined based on the loci-IF interaction map shown in Figure 2c. Randomized data was generated by scrambling the cluster identities of individual loci in cells while keeping the total number of loci within each cluster the same within that cell. The contact probability for observed and randomized data for each cell are shown as boxplots. b, In individual cells, loci associated with each cluster is mapped onto the chromosome structure images shown in Fig 1g. c, Histogram of IF marks for each loci in each of the clusters. Cluster 1 is enriched in repressive markers such as H3K9me3, mH2A1, DAPI. Cluster 2 is enriched in interactions with Fibrillarin and is associated with nucleolus. Cluster 3 is enriched in active marks such as RNAPII Ser5-P, H3K27ac and SF3a66 (nuclear speckle marker). Cluster 4 is enriched in Lamin B1 and is associated with nuclear lamina. d, Mapping of the A/B compartment definitions^4^ onto the tSNE plot based on the loci-IF mark interaction map shown in Figure 2c. Note that regions that are not assigned to one of the compartments in the study were excluded from the analysis. (n = 1,188, 960 loci in A and B compartment). e, Histogram of the IF marks for each loci assigned to A or B compartments. f, The contact probability between loci in A/B compartments similar to (a). For the boxplots in a, f, the center line in the boxes marks median, the upper and lower limits of the boxes mark the interquartile range, the whiskers extend to the farthest data points within 1.5 times the interquartile range, and the gray points mark outliers. g, Images of individual cells with loci associated with A/B mapped onto the chromosome, similar to Fig 2g. Observed compared to shuffled data for 2 cells, shown in Fig 2g and (b) respectively. h, The full contact map between all loci from the 1Mb DNA seqFISH+ data with a search radius of 1 μm. i, In single cells, the chromosome to chromosome contact maps considering only the first 5 Mb regions with a search radius of 2.5 μm. j, Similar to i, but considering only the last 5 Mb of the chromosome. Showing that single cell heterogeneity in the contacts between the proximal and distal regions of the chromosomes, close to the repetitive regions. n = 201 cells in a, c, d-f, h.

**Extended Data Fig. 7.**
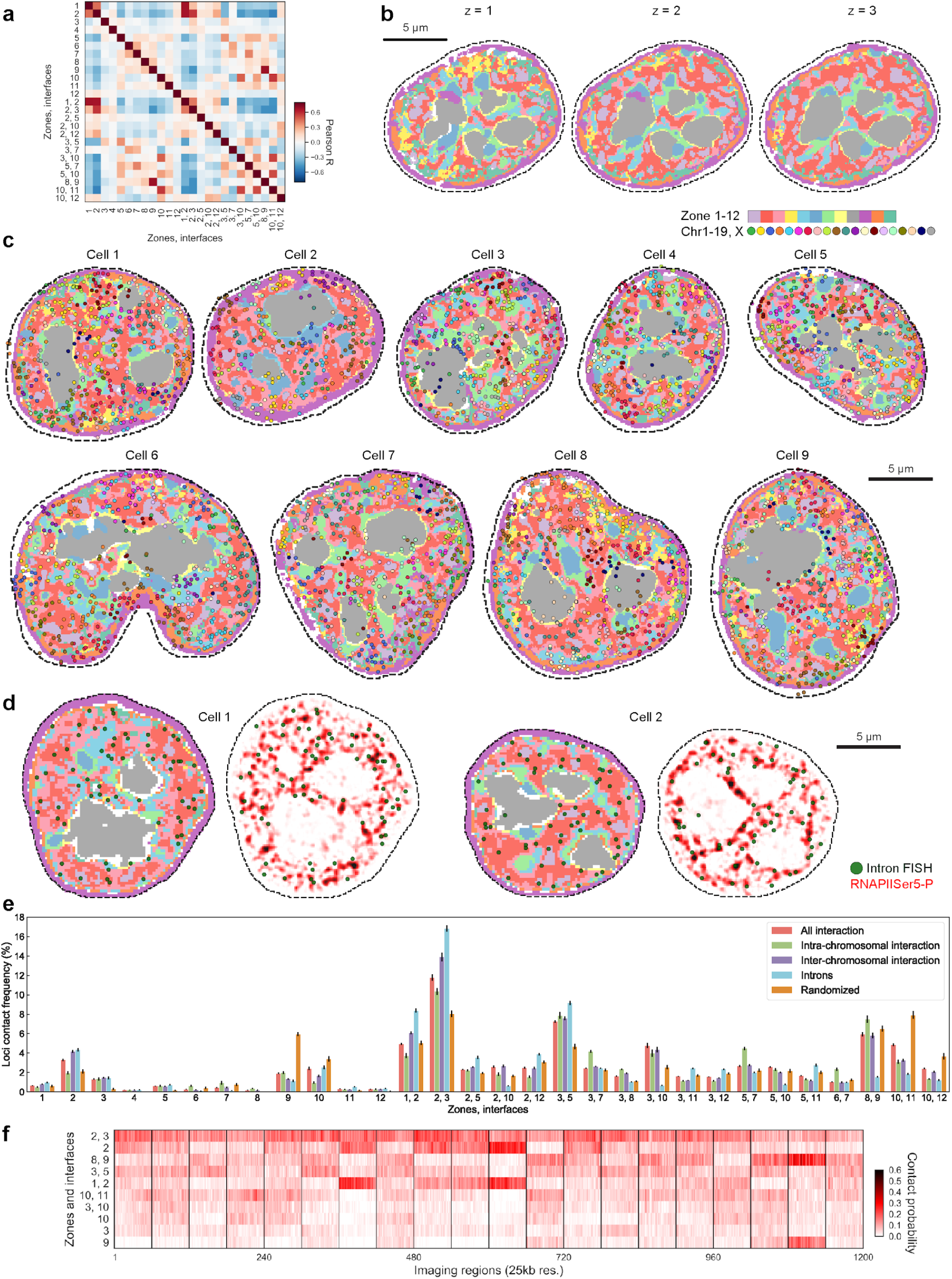
Additional visualization and quantification for combinatorial subcellular chromatin patterns or nuclear zones. a, Correlation matrix between zones and interfaces based on the DNA loci association with zones and interfaces shown in Figure 3e (n = 2,460 loci). Loci appearing in zone 1 are also more likely to be found in zone 2 as well as in interface 1, 2. b, Reconstruction of zones in the cell shown in Fig 3d with additional z-plane. c, Reconstruction of zones and DNA loci in additional cells. d, Reconstruction of zones and 1,000 gene intron dots as well as RNAPIISer5-P staining (background-subtracted) in additional cells. c, Contact frequency between DNA loci and zones/interfaces in single cells, calculated for all loci, loci contacting other loci from the same chromosomes (intra-chromosomal), interchromosomal, transcriptional active sites measured by intron FISH, and random loci (randomized control). d, Heatmap for contact probability between DNA loci, nuclear zones and interfaces for the 25 kb data. Loci within the same Mb region have similar nuclear zone and interface contact probability. (n = 201 cells in a-f).

**Extended Data Fig. 8.**
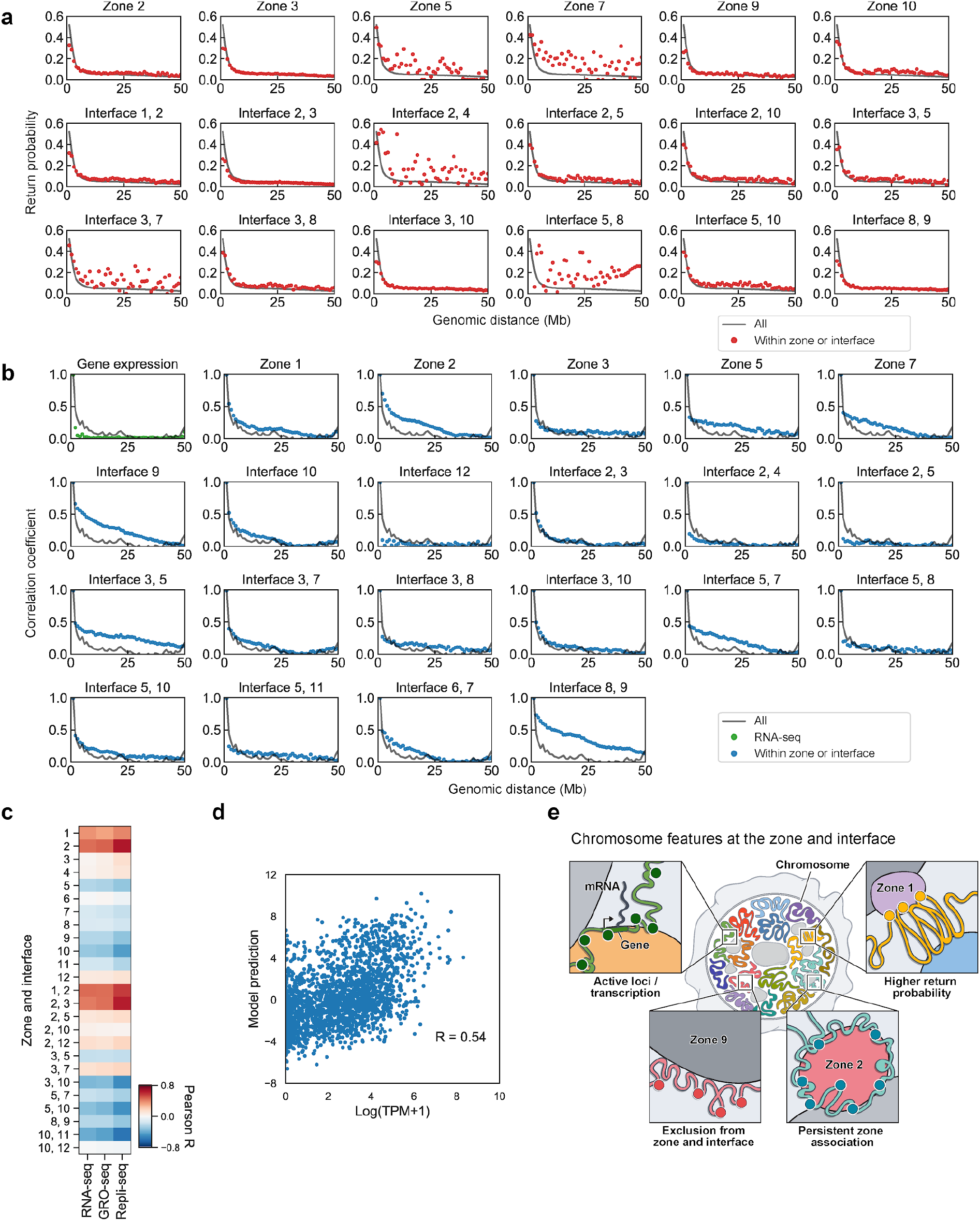
Further characterization of nuclear zones and interfaces. a, Auto-correlation function of the frequency of loci contacting a particular zone (blue) or correlated across all zones (black). Gene expression correlation across loci are shown in green. Expression levels of loci decorrelate rapidly within 1 Mb, while zone association could decay more slowly over several Mbs. b, Return probability as a function of genomic distance, showing higher return probability for loci within some zone or interface (red) compared to all loci (black). c, Correlation between zone association and gene expression levels (RNA-seq)^34^, density of RNA polymerases on the loci (GRO-seq)^46^ and early replication domains (Repli-seq)^47^ for all loci (n = 2,460 loci). d, A linear model (see ‘Methods’) to estimate expression level based on the zone association for each loci (each dot) is compared against the expression levels measured by RNA-seq^34^. (n = 2,460 loci) R represents Pearson correlation coefficient. (n = 201 cells in a-d). e, An Illustration of the nuclear features observed in the DNA seqFISH+ dataset.

**Extended Data Fig. 9.**
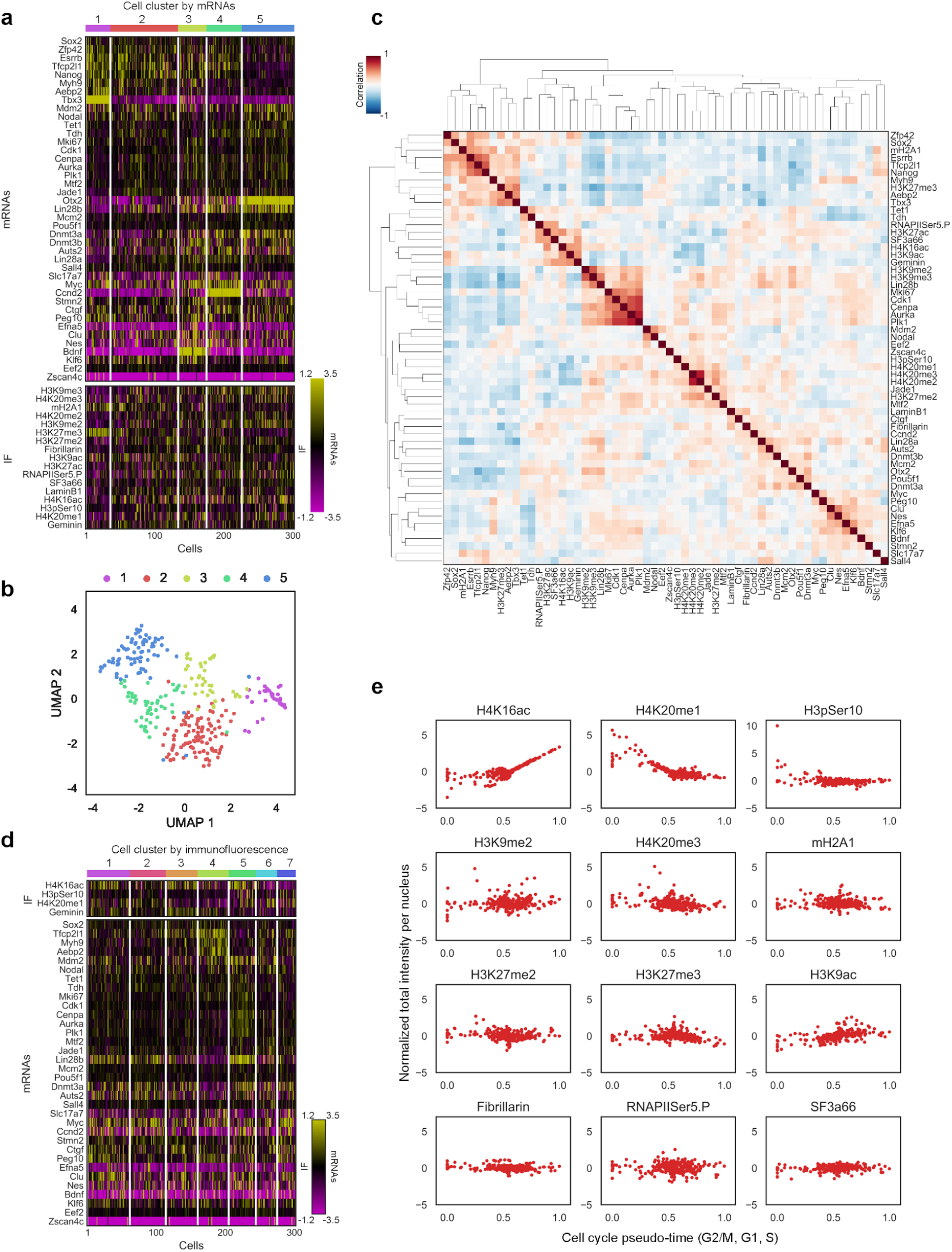
Heterogeneity of transcriptional and chromatin states and their relationships in single cells. a, Heatmap of cell clusters with distinct mRNA profiles, showing clusters (1, 2) expressing pluripotency markers (e.g., Nanog, Esrrb, Zfp42) and close to ground state of pluripotency, clusters (3, 4) that are primed for differentiation while remaining pluripotency factor expression and a cluster (5) more close to differentiation, consistent with single cell RNA-seq studies in mESCs^30–34^. IF marks are shown with cells within each cluster. b, UMAP representations of the cell clusters defined by mRNA profiles. c, Joint correlation matrix between mRNA and IF markers based on the normalized count or intensity profiles in single cells. Color bar represents Pearson’s correlation coefficient. (n = 58 mRNA and IF markers) d, Heatmap of cell clusters with distinct IF profiles shown with cell cycle associated IF markers and additional mRNA markers, similar to Fig 4b. Cell cycle associated IF markers were excluded from clustering steps and do not show strong enrichments in the IF clusters. e, Pseudo-time course analysis for cell cycle progression, cell cycle markers (top) show clear enrichments while other markers (middle and bottom panels) do not show specific enrichments upon cell cycle pseudo-time course, suggesting majority of the IF markers profiled are not primarily affected by cell cycle phases. n = 302 cells in the center field of views from two replicates in a-e.

**Extended Data Fig. 10.**
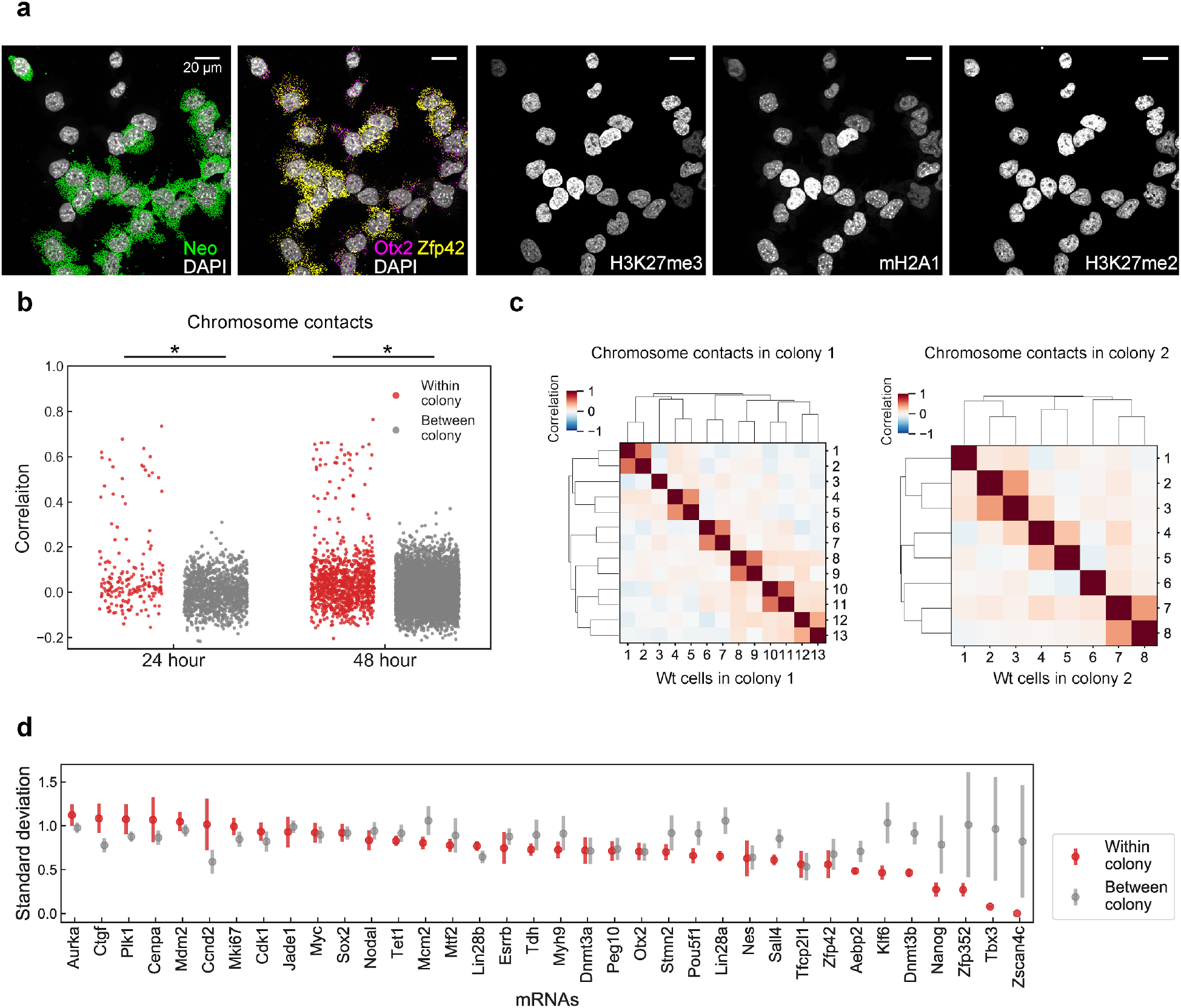
Additional analysis for colony level cell state heterogeneity. a, mRNAs and IF images in a colony in the 48 hour clonal tracing experiment. H3K27me3 and mH2A1 overall intensities are similar in WT cells (GFP/Neo negative) in the colony. b, Histogram of cell-to-cell correlations of chromosome to chromosome contacts maps for cells within colonies (red) and between colonies (grey). Cells with similar chromosome structures (red dots with high correlation values) are likely to be sister cells. Y-axis represents Pearson’s correlation coefficient, computed by 20 x 20 chromosome contact matrices from pairs of cells. *p < 0.001 (p = 2.2e-9, 7.2e-21) from a two-sided Wilcoxon’s rank sum test with pairs of cells of 180, 1,198, 966 and 5,820 (from left to right). c, Correlation of chromosome contacts between cells in colonies in the 48 hour clonal tracing experiment. Strong correlations are seen between putative sister cells suggesting that gross chromosome structures are preserved for 1 generation. Color bars represent Pearson’s correlation coefficient. d, Standard deviation of normalized mRNA levels within colonies (red) and between colonies (grey). Tbx3 and Nanog are more homogeneous within colonies, consistent with previous findings of long-lived Nanog states across several generations based on single cell live imaging experiments^31,34^. n = 117 unlabeled cells within colonies from a 48-hour dataset.

**Supplementary Table 1** List for genomic coordinates of the 3,660 DNA loci used in DNA seqFISH+.

**Supplementary Table 2** Codebook for the 3,660 DNA loci in the three fluorescent channels. Base 16 pseudocolor coding scheme for each of the loci in the channel 1 and 2, include control regions for off target evaluation. Region and chromosome paint imaging scheme for each of the loci in the channel 3.

**Supplementary Table 3** Chromatin profiles for each antibody and genomic locus.

**Supplementary Table 4** Normalized association frequencies of individual zones and interfaces on each of the 3,660 DNA loci.

**Supplementary Table 5** List for target RNAs in E14 replicates and clonal experiments.

## Methods

### Data reporting

No statistical methods were used to predetermine sample size. The experiments were not randomized and the investigators were not blinded to allocation during experiments and outcome assessment.

### DNA seqFISH+ encoding strategy

A 16-base coding scheme with 5 rounds of barcoding is used in DNA seqFISH+ for the 1 Mb resolution data in fluorescent channel 1 (643-nm) and 2 (561-nm) (Supplementary Table 2). The first 3 rounds of barcoding codes for 16^3=4,096 unique barcodes. Two additional rounds of parity check (linear combinations of the first three rounds) are included. 2,048 barcodes are selected to correct for dropouts in any 2 out of 5 rounds of barcoding and used in both channel 1 and 2. The 16-pseudocolor base is generated by hybridizing the sample with 16 different readout oligos sequentially.

To image 20 distinct regions (1.5-2.4 Mb) with 25 kb resolution, a combined strategy of diffraction limited spot imaging and chromosome painting is used in channel 3 (488-nm) (Supplementary Table 2). For the initial 60 rounds, 25 kb regions are readout one at a time on all 20 chromosomes in each round of hybridization. These 60 rounds can resolve the 25kb loci within each distinct region but cannot distinguish which chromosome the loci belong to. The next 20 rounds are used to resolve the identities of the 20 distinct regions or chromosomes by painting the entire region (1.5-2.4 Mb) one at one time. With this strategy, identities for 1,200 loci are decoded.

In total, 80 readouts are used in each fluorescent channel for a total of 240 readouts for 3 channels.

### Primary probe design

RNA seqFISH probes were designed as described previously^27,28^. In brief, 35-nt RNA target binding sequences, 15-nt unique readout probe binding sites for each RNA target, and a pair of 20-nt primer binding sites at 5’ and 3’ end of the probe for probe generation are concatenated. (see ‘primary probe synthesis’). Marker genes (Supplementary Table 5) were selected based on previous single cell imaging and RNA-seq studies in mESCs^27,31–34^.

For DNA seqFISH+ target region selection (Supplementary Table 1), the unmasked and repeat-masked GRCm38/mm10 mouse genome FASTA files were downloaded from Ensembl release 93^55^. To select target regions for channel 1, the entire mouse genome was split into candidate target regions of 25 kb. Masking coverage was evaluated for each region using the repeat-masked genome. Regions with a high percentage of masked bases were removed from consideration. Then target regions were further selected to space out approximately 2 Mb in the genome coordinates. To select target regions for channel 2, candidate genes related to mESCs pluripotency and differentiation were selected from previous studies^7,32,33^, and then 25 kb regions were selected by centering the transcription starting sites of the genes. To select target regions for channel 3, gene loci with various expression levels in mESCs as well as gene poor regions were initially selected as a 2.5 Mb block, and splitted into 25 kb blocks. Only a single 2.5 Mb region was selected per chromosome.

Region-specific primary probes were designed as previously described for single-stranded RNA^28^ with some modifications. The target region was extracted from the unmasked genome. Probe sequences were produced by taking the reverse complement of 35-nt sections of the target region. Starting from the 5’ end of the forward strand, candidate probes were tested for viability, shifting one base at a time. Probes that contained five or more consecutive bases of the same kind, or had a GC content outside of 45-65%, were considered non-viable. Each time a viable probe was discovered, evaluation was switched to the opposite strand, starting 19-nt downstream from the start of the viable probe to mitigate cross-hybridization between neighboring probes. This procedure was repeated until the end of the target region was reached.

Next, the probes were aligned to the unmasked mouse genome for off-target evaluation using Bowtie2^56^. Any alignment containing at least 19 matched bases that fell outside the genomic coordinates of the target region was considered off-target. Probes with more than 10 total off-target hits were dropped. Off-target hits were grouped into 100 kb bins and stored for use in the final probe selection. Bins were overlapped by 50 kb so that closely grouped hits could not evade the filter by splitting into two bins. Additionally, probes were checked for matches with a BLAST^57^ database constructed from common repeating sequences in mammals. The FASTA file for “Simple Repeat” sequences for “Mammalia only” was downloaded from Repbase^58^. All probes with at least 19 matched bases with the repeats index were dropped. After filtering the probes, all remaining probes were evaluated for potential cross-hybridization using BLAST^57^. Any probe pairs with at least 19 matched bases were dropped in the final probe selection.

Final probe sets were selected to maintain probe specificity, and to achieve a relatively uniform spacing of probes on the target sequence. Final probes were selected one by one, starting with the target region with the fewest remaining probes. The probe that minimized the sum of the squares of distances between adjacent selected probes and the start and end coordinates of the target region was selected. After selecting a probe, any probes that were found to cross-hybridize with at least 19-nt to the selected probe were dropped. As probes were added, their off-target hits were summed by bin. If the addition of a probe resulted in any bin having 10 total hits, all remaining unselected probes that had an off-target hit in that bin were dropped. For channel 1 and 2 probes, once 200 probes were selected for a target region, all remaining probes for that region were dropped. For the channel 3 probes, regions containing up to 150 probes were kept and other regions were dropped, and as a result, 1.5-2.4 Mb of 20 distinct regions containing 60 of 25 kb regions were finally selected.

Primary probes were then assembled similar to previous seqFISH studies^26–28,59,60^. For Mb resolution DNA seqFISH+ in channel 1 and 2, primary probes consist of the genomic region specific 35-nt sequences, flanked by the five unique 15-nt readout probe binding sequences, which correspond to pseudo-channel in each barcoding round, and a pair of 20-nt primer binding sites at the 5’ and 3’ end of the probe. For 25 kb resolution DNA seqFISH+ in channel 3, primary probes consist of the genomic region specific 35-nt sequences, flanked by three identical binding sites of a 15-nt readout probe, which corresponds to one of the 60 sequential rounds for the diffraction limited spot imaging, and two identical binding sites for a 15-nt readout probe, which corresponds to one of the 20 distinct regions for the chromosome painting, and 20-nt primer binding sites at the 5’ and 3’ end of the probes.

### Primary probe synthesis

Primary probes were generated from oligoarray pools (Twist Bioscience) as previously described^26–28,59,60^ with some modifications. In brief, probe sequences were amplified from the oligo pools with limited two-step PCR cycles (first step PCR primers, 4-fwd: 5’-ATGCGCTGCAACTGAGACCG; 4-rev: 5’-CTCGACCAAGGCTGGCACAA; second step PCR primers, 4-fwd: 5’-ATGCGCTGCAACTGAGACCG; 4-T7rev: 5’-TAATACGACTCACTATAGCTCGACCAAGGCTGGCACAA)^26–28,59,60^, and PCR products were purified using QIAquick PCR Purification Kit (Qiagen 28104). Then in vitro transcription (NEB E2040S) followed by reverse transcription (Thermo Fisher EP7051) were performed. For the DNA seqFISH+ primary probes, the forward primer (4-fwd) with 5’ phosphorylation was used to allow ligation of the primary probes as described below (see ‘Cell culture experiment’). After reverse transcription, the single-stranded DNA (ssDNA) probes were alkaline hydrolysed with 1 M NaOH at 65°C for 15 min to degrade the RNA templates, and then neutralized with 1 M acetic acid. Then, probes were ethanol precipitated, and eluted in nuclease-free water.

LINE1 and SINEB1 probes were similarly generated except using mouse genomic DNA template extracted from E14 mESCs with DNeasy Blood & Tissue Kits (Qiagen 69504) for PCR. Primers for LINE1 and SINEB1^35^ contain readout probe binding sites as overhangs to allow readout probe hybridization and stripping with seqFISH routines. Genome targeting sequences of the primary probes were 113-nt and 117-nt for LINE1 and SINEB1, respectively.

### Readout probe design and synthesis

Readout probes of 12-15 nucleotides in length were designed for seqFISH as previously described^27,28^. In brief, a set of probe sequences was randomly generated with combinations of A, T, G or C nucleotides with a GC-content range of 40-60%. To minimize cross-hybridization between the readout probes, any probes with ten or more contiguously matching sequences between the readout probes were removed. The readout probes for sequential immunofluorescence were similarly designed except ‘C’ nucleotide is omitted^61^. The 5’ amine-modified DNA oligonucleotides (Integrated DNA Technologies) with the readout probe sequences were conjugated to Alexa Fluor 647-NHS ester (Invitrogen A20006) or Cy3B-NHS ester (GE Healthcare PA63101) or Alexa Fluor 488-NHS (Invitrogen A20000) as described before^27,28^, or fluorophore conjugated DNA oligonucleotides were purchased from Integrated DNA Technologies.

### DNA-antibody conjugation

Preparation of oligo DNA conjugated primary antibodies was performed as described before^62^ with modifications. In brief, to crosslink thiol-modified oligonucleotides to lysine residues on antibodies, BSA-free antibodies were purchased from commercial vendors whenever possible. Antibodies (90-100 μg) were buffer-exchanged to 1× PBS using 7K MWCO Zeba Spin Desalting Columns (Thermo Scientific 89882), and reacted with 10 equivalent of PEGylated SMCC cross-linker (SM(PEG)2) (Thermo Scientific 22102) diluted in anhydrous DMF (Vector Laboratories S4001005). The solution was incubated at 4°C for 3 hours, and then purified using 7K MWCO Zeba Spin Desalting Columns. In parallel, 300 μM 5’ thiol-modified 18-nt DNA oligonucleotides (IDT) were reduced by 50 mM dithiothreitol in 1× PBS at room temperature for 2 hours, and purified using NAP5 columns (GE Healthcare 17-0853-01). Then maleimide activated antibodies were mixed with 6-15 equivalent of the reduced form of the thiol-modified DNA oligonucleotides in 1× PBS at 4°C overnight. DNA-primary antibody conjugates were washed with 1× PBS four times and concentrated using 50 KDa Amicon Ultra Centrifugal Filters (Millipore, UFC505096). The concentration of conjugated oligo DNA and antibody with BCA Protein Assay Kit (Thermo Scientific 23225) were quantified using Nanodrop.

For the BSA containing primary antibodies, SiteClick R-PE Antibody Labeling Kit (Life Technologies S10467) was used to conjugate the antibodies with 10-20 equivalent of 5’ DBCO-modified 18-nt DNA oligonucleotides (IDT). The oligo conjugated antibodies were validated by SDS-PAGE gel and imaging, and stored in 1x PBS at −80°C as small aliquots.

### Cell culture and preparation

E14 mESCs (E14Tg2a.4) from Mutant Mouse Regional Resource Centers were maintained under serum/LIF condition as previously described^27,31^. A stable E14 line that targets endogenous repetitive regions with the CRISPR/Cas system^63^ was generated similarly to the previous study^26^. In brief, PiggyBac vectors, PGK-NLS-dCas9-NLS-3xEGFP, carrying a separate puromycin resistance cassette under an EF1 promoter, and mU6-sg3632454L22Rik(F+E), carrying a separate neomycin resistance cassette under a SV40 promoter, were constructed. A single-guide RNA (sgRNA) sequence (5’-GGAAGCCAGCTGT) was used to target repetitive regions at the 3632454L22Rik gene locus in X chromosome. To create the stable E14 line (GFP/Neo E14) with those vectors, transfection was performed with FuGENE HD Transfection Reagent (Promega E2311), and cells were selected with puromycin (Gibco A1113803) at 1 μg/mL. After the selection, single clones were isolated manually, and stable labeling of the locus was verified by imaging.

E14 cells were plated on poly-D-lysine (Sigma P6407) and human laminin (BioLamina LN511) coated coverslips (25 mm x 60 mm)^27^, and incubated for 24 or 48 hours. Then cells were fixed with freshly made 4% formaldehyde (Thermo Scientific 28908) in 1× PBS (Invitrogen AM9624) at room temperature for 10 minutes. The fixed cells were washed with 1× PBS a few times, and stored in 70% ethanol at −20°C. In the case of co-culture experiments with unlabeled E14 cells and the GFP/Neo E14 cells (monoclonal line), cell densities were counted and cell lines were mixed with a 1:10 ratio.

### Cell culture experiment

The fixed and stored cell samples were dried, and permeabilized with 0.5% Triton-X (Sigma-Aldrich 93443) in 1× PBS at room temperature for 15 minutes after attaching a sterilized silicon plate (McMASTER-CARR 86915K16) with an punched hole to the coverslip to use it as a chamber. The samples were washed three times with 1× PBS and blocked at room temperature for 15 minutes with blocking solution consisted of 1× PBS, 10 mg/mL UltraPure BSA (Invitrogen AM2616), 0.3% Triton-X, 0.1% dextran sulfate (Sigma D4911) and 0.5 mg/mL sheared Salmon Sperm DNA (Invitrogen AM9680). Then DNA oligo-conjugated primary antibodies listed below were incubated in the blocking solution with 100-fold diluted SUPERase In RNase Inhibitor (Invitrogen AM2694) at 4°C overnight. The typical final concentration of DNA conjugated primary antibodies used were estimated as 1-5 ng/μL. The samples were washed with 1× PBS three times and incubated at room temperature for 15 minutes, before postfixing with freshly made 4% formaldehyde in 1× PBS at room temperature for 5 minutes. Next, the samples were washed with 1× PBS six times and incubated at room temperature for 15 minutes. The samples were then further post-fixed with 1.5 mM BS(PEG)5 (PEGylated bis(sulfosuccinimidyl)suberate) (Thermo Scientific A35396) in 1× PBS at room temperature for 20 minutes, followed by quenching with 100 mM Tris-HCl pH7.4 (Alfa Aesar J62848) at room temperature for 5 minutes. After the post-fixation, the samples were washed with 1xPBS and air dried after removing the custom silicon chamber.

The oligo DNA conjugated primary antibodies used were as follows: mH2A1 (Abcam ab232602), E-Cadherin (R&D AF748), Fibrillarin (C13C3) (Cell Signaling 2639BF), Geminin (Abcam ab238988), GFP (Invitrogen G10362), H3 (Active Motif 39763), H3K27ac (Active Motif 39133), H3K27me2 (Cell Signaling 9728BF), H3K27me3 (Cell Signaling 9733BF), H3K4me1 (Cell Signaling 5326S), H3K4me2 (Cell Signaling 9725BF), H3K4me3 (Active Motif 39915), H3K9ac (Active Motif 91103), H3K9me2 (Abcam ab1220), H3K9me3 (Diagenode MAb-146-050), H3pSer10 (Millipore 05-806), H4K16ac (EMD Millipore 07-329), H4K20me1 (Abcam ab9051), H4K20me2 (Abcam ab9052), H4K20me3 (Active Motif 39671), Lamin B1 (Abcam ab220797), RNAPII Ser5-P (Abcam ab5408), SF3a66 (Abcam ab77800). Two antibodies (E-Cadherin and GFP) were only included in the clonal tracing experiments. Several antibodies (H3, H3K4me1, H3K4me2 and H3K4me3) were excluded from the downstream analysis due to the quality of antibody staining with oligo-conjugation.

After the immunofluorescence preparation above, custom-made flow cells (fluidic volume ~30 μl), which were made from glass slide (25 x 75 mm) with 1 mm thickness and 1 mm diameter holes and a PET film coated on both sides with an acrylic adhesive with total thickness 0.25 mm (Grace Bio-Labs RD481902), were attached to the coverslips. The samples were rinsed with 2× SSC, and RNA seqFISH primary probe pools (1-10 nM per probe) and 10 nM polyT LNA oligo with a readout probe binding DNA sequence (Qiagen) were hybridized in 50% hybridization buffer consisted of 50% formamide (Invitrogen AM9342), 2× SSC and 10% (w/v) dextran sulfate (Millipore 3710-OP). The hybridization was performed at 37°C for 24-72 hours in a humid chamber. After hybridization, the samples were washed with a 55% wash buffer consisting of 55% formamide, 2× SSC and 0.1% Triton X-100 at room temperature for 30 minutes, followed by three rinses with 4× SSC. Then samples were imaged for RNA seqFISH as described below (see ‘seqFISH imaging’). Note that immunofluorescence signals can be imaged at this step.

After RNA seqFISH imaging, the samples were processed for DNA seqFISH+ primary probe hybridization. The samples were rinsed with 1× PBS, and incubated with 100-fold diluted RNase A/T1 Mix (Thermo Fisher EN0551) in 1× PBS at 37°C for 1 hour. Then samples were rinsed three times with 1× PBS, followed by three rinses with a 50% denaturation buffer consisting of 50% formamide and 2× SSC and incubation at room temperature for 15 minutes. Then the samples were heated on the heat block at 90°C for 4.5 minutes in the 50% denaturation buffer, by sealing the inlet and outlet of the custom chamber with aluminum sealing tapes (Thermo Scientific 232698). After heating, the samples were rinsed with 2× SSC, and DNA seqFISH+ primary hybridization buffer consisting of ~1 nM per probe, ~1 μM LINE1 probe, ~1 μM SINEB1 probe, 100 nM 3632454L22Rik fiducial marker probe (IDT), 40% formamide, 2× SSC and 10% (w/v) dextran sulfate (Millipore 3710-OP) was hybridized at 37°C for 48-96 hours in a humid chamber. After hybridization, the samples were washed with a 40% wash buffer consisting of 40% formamide, 2× SSC and 0.1% Triton X-100 at room temperature for 15 minutes, followed by three rinses with 4× SSC.

Then samples were further processed to “padlock” primary probes. A global ligation bridge oligo (IDT) was hybridized in a 20% hybridization buffer consisting of 20% formamide, dextran sulfate (Sigma D4911) and 4xSSC at 37°C for 2 hours. The 31-nt global ligation bridge (5’-TCAGTTGCAGCGCATGCTCGACCAAGGCTGG) was designed to hybridize to 15-nt of the DNA seqFISH+ primary probes at 5’ end and 16-nt at the 3’ end. Then, samples were washed with 10% WB for three times and incubated at room temperature for 5 minutes. After three rinses with 1× PBS, the samples were then incubated with 20-fold diluted Quick Ligase in 1× Quick Ligase Reaction Buffer from Quick Ligation Kit (NEB M2200) supplemented with additional 1 mM ATP (NEB P0756) at room temperature for 1 hour to allow ligation reaction between 5’- and 3’-end of the DNA seqFISH+ primary probes. Then the samples were washed with a 12.5% wash buffer consisting of 12.5% formamide, 2× SSC and 0.1% Triton X-100, followed by three rinses with 1× PBS.

The samples were then processed for amine modification and post-fixation to further stabilize the primary probes. The samples were rinsed with 1× Labeling Buffer A, followed by incubation with 10fold diluted Label IT Amine Modifying Reagent in 1× Labeling Buffer A from Label IT Nucleic Acid Modifying Reagent (Mirus Bio MIR 3900) at room temperature for 45 minutes. After three rinses with 1× PBS, the samples were fixed with 1.5 mM BS(PEG)5 in 1× PBS at room temperature for 30 minutes, followed by quenching with 100 mM Tris-HCl pH7.4 at room temperature for 5 minutes. The samples were washed with a 55% wash buffer at room temperature for 5 minutes, and rinsed with 4× SSC for three times. Then samples were imaged for DNA seqFISH+ and sequential immunofluorescence as described below (see ‘seqFISH imaging’).

The 1,000 gene intron experiments in Fig. 3d, e and Extended Data Fig. 7d, e were performed similarly with minor modifications. E14 coverslips were prepared and processed by following the sequential immunofluorescence steps above. After the sequential immunofluorescence preparation, 1,000 gene intron FISH probes^27^were hybridized in the 50% hybridization buffer at 37°C for 24 hours in a humid chamber. Then samples were washed with the 55% wash buffer at 37°C for 30 minutes, followed by three rinses with 4× SSC. Then samples were imaged for intron FISH and sequential immunofluorescence as described below (see ‘seqFISH imaging’).

### Microscope setup

All imaging experiments were performed with the imaging platform and fluidics delivery system similar to those previously described^27,28^. The microscope (Leica DMi8) was equipped with a confocal scanner unit (Yokogawa CSU-W1), a sCMOS camera (Andor Zyla 4.2 Plus), 63× oil objective lens (Leica 1.40 NA), and a motorized stage (ASI MS2000). Fiber coupled lasers (643, 561, 488 and 405 nm) from CNI and Shanghai Dream Lasers Technology and filter sets from Semrock were used. The custom-made automated sampler was used to move designated readout probes in hybridization buffer from a 2.0 mL 96-well plate through a multichannel fluidic valve (IDEX Health & Science EZ1213-820-4) to the custom-made flow cell using a syringe pump (Hamilton Company 63133-01). Other buffers were also moved through the multichannel fluidic valve to the custom-made flow cell using the syringe pump. The integration of imaging and the automated fluidics delivery system was controlled by custom written scripts in μManager^64^.

### seqFISH imaging

The sequential hybridization and imaging routines were performed similarly to those previously described^27,28^ with some modifications. In brief, the sample with the custom-made flow cell was first connected to the automated fluidics system on the motorized stage on the microscope. Then the regions of interest (ROIs) were registered using nuclei signals stained with 5 μg/mL DAPI (Sigma D8417) in 4× SSC. RNA seqFISH imaging was performed with the sequential hybridization and imaging routines described below first. After the completion of RNA seqFISH imaging, the samples were disconnected from the microscope, and proceeded to the DNA seqFISH+ procedures (see ‘Cell culture experiment’). For the DNA seqFISH+ and sequential IF imaging, the registered ROIs for RNA seqFISH were loaded and manually corrected to ensure to image the same ROIs as RNA seqFISH imaging, and following routines were performed.

All the sequential hybridization and imaging routines below were performed at room temperature. The serial hybridization buffer contained two or three unique readout probes (10-25 nM) with different fluorophores (Alexa Fluor 647, Cy3B or Alexa Fluor 488) in 10% EC buffer (10% ethylene carbonate (Sigma E26258), 10% dextran sulfate (Sigma D4911) and 4× SSC), and was picked up from a 96-well plate and flow into the flow cell for 20 minutes incubation. For DNA seqFISH+ experiments, readout probes (Alexa Fluor 647, Cy3B or Alexa Fluor 488) for sequences designated as fiducial markers were also included in the serial hybridization buffer to allow image registration at the subpixel resolution. After the serial hybridization, the samples were washed with 1 mL of 4× SSCT (4× SSC and 0.1% Triton-X), followed by a wash with 330 uL of the 12.5% wash buffer. Then, the samples were rinsed with ~200 μl of 4× SSC, and stained with ~200 uL of the DAPI solution for 30 s. Next, anti-bleaching buffer was flown through the sample for imaging. The anti-bleaching buffer was made of 50 mM Tris-HCl pH 8.0 (Invitrogen 15568025), 300 mM NaCl (Invitrogen AM9759), 2× SSC, 3 mM trolox (Sigma 238813), 0.8% D-glucose (Sigma G7528), 1,000-fold diluted catalase (Sigma C3155), 0.5 mg/mL glucose oxidase (Sigma G2133)^27^ for E14 experiments, and made of 50 mM Tris-HCl pH 8.0, 4× SSC, 3 mM trolox, 10% D-glucose, 100-fold diluted catalase, 1 mg/mL glucose oxidase (Sigma G2133)^28^ for unlabeled E14 and GFP/Neo E14 line clonal experiments.

Snapshots were acquired with 0.25 μm z-steps over 6 μm z-slices with 643-nm, 561-nm, 488-nm and 405-nm fluorescent channels per field of view, except for RNA seqFISH in the clonal experiments acquired with 0.75 μm z-steps with 643-nm, 561-nm, 488-nm fluorescent channels. After image acquisition, 1 mL of the 55% wash buffer was flown for 1 minutes to strip off readout probes, followed by an incubation for 1 minutes before rinsing with 4× SSC. The serial hybridization, imaging and signal extinguishing steps were repeated until the completion of all rounds. During the RNA seqFISH and DNA seqFISH+ imaging routines, blank images containing only autofluorescence of the cells were imaged at the beginning and end of the routines. During the DNA seqFISH+ imaging, images containing only fiducial markers were also imaged at the beginning and at the end of the routines for the image alignment (see ‘Image Analysis’). Images were manually checked at the end of all imaging routines and in case problematic hybridization rounds such as off-focus appeared, those hybridization rounds were repeated.

### Image Analysis

To correct for the non-uniform background, a flat field correction was applied by dividing the normalized background illumination with each of the fluorescence images while preserving the intensity profile of the fluorescent points. The background signal was then subtracted using the ImageJ rolling ball background subtraction algorithm with a radius of 3 pixels.

FISH spot locations were obtained by using a laplacian of gaussians filter, semi-manual thresholding as described below, and a 3D local maxima finder. Subsequently the locations were super resolved using a 3D radial center algorithm^65,66^. Briefly, a 3×3×3 cube of pixels around a local maxima found above the specified threshold was taken from the aligned and background subtracted image. This sub-image was then used to calculate the sub-pixel location of the RNA molecule or DNA locus and the mean standard deviation (average of the standard deviation in each dimension) of the intensity cloud using a 3D radial center algorithm. A MATLAB implementation of the algorithm can be found on the Parthasarathy lab website. The resulting RNA or DNA spot locations were further filtered based on the size of the sigma values.

To find the optimal threshold values for the spot detection, threshold values for RNA seqFISH were updated manually. In contrast, for DNA seqFISH+, 29 incremental threshold values, were initially applied to the images in the first position. The number of spots and median spot intensity in the nuclei were computed for each of the 29 thresholds across 80 hybridizations. Then the threshold value for the first hybridization round was manually chosen, and threshold for the other hybridizations were selected such that the number of dots detected matches most closely to those expected from the codebook. For example, if hyb 1 targets 30 loci and hyb 2 targets 60 loci, then hyb 2 should have twice as many dots as hyb 1. In this process, we assumed all loci can be detected with the same detection efficiency on average. In addition, the median intensities from the adjacent threshold values were compared, and whenever intensity differences are more than 15%, a more stringent threshold value was taken to fulfill this criteria to minimize non-specific spot detection. These processes were performed in individual fluorescent channels independently. Similarly, we corrected the threshold values across positions by computing the ratio of the median intensities relative to those from the first position per hybridization in order to minimize detection bias across different positions.

To align spots or images in different channels to those in the reference channel (643-nm), chromatic aberration shifts were corrected using the fiducial markers to calculate the offsets. To align RNA seqFISH and sequential immunofluorescence images in different hybridization rounds, reference channels (either DAPI or polyA staining) were aligned using 2D phase correlations along every axis iteratively to find a consensus transformation for alignment as described before^27^. The 2D phase correlation algorithm is implemented in MATLAB with the function imregcorr. To align DNA seqFISH+ spots in different hybridization rounds, fiducial markers were identified in each image by searching for the known ‘constellation’ seen in images containing only the fiducial markers. To identify a first pair of distant fiducial markers, the vector describing the relative position of the known markers was compared with those separating similarly oriented pairs of FISH spots in each image. Most, if not all, of the fiducial marker ‘constellation’ can then be recovered by searching for each fiducial marker at its known location relative to that of previously identified fiducial markers in the image. Further alignment to correct any rotation between RNA and DNA FISH images was done as follows. First, both image stacks to be aligned (DAPI staining) were converted to 2D images using a maximum intensity projection in the z-dimension. The resulting 2D images were aligned using a one plus one evolutionary optimization method to maximize the Mattes Mutual Information between the images with the transformation constrained to only rigid transforms with a maximum of 500 iterations. This algorithm is implemented in MATLAB with the function imregtform. Once 2D alignment with both translation and rotation was obtained, one stack was transformed using the found transformation. The image stacks were then projected along the x axis and aligned using a normalized cross-correlation to determine the first estimate of the z-dimension offset. The image was then projected along the y axis to find a second estimate of the z-dimension offset using the same method. The two offsets were averaged.

To assign mRNA spots to individual cells, the processed spots were collected within individual cytoplasmic ROIs, which were segmented manually from polyA or E-Cadherin images. Similarly, to assign intron spots to individual cells, intron spots within individual nuclear ROIs from DAPI images^27^ were collected. By comparing the centroids between cytoplasmic ROIs and nuclear ROIs, numbers from both ROIs were matched. Only cells at the center of the fields of view were preserved for the RNA analysis to avoid biasing the RNA distribution.

For channel 1 and 2 barcode decoding in DNA seqFISH+, once all potential points in all hybridizations were obtained, points were matched to potential barcode partners in all other barcoding rounds of all other hybridizations using a 1.73 pixel search radius to find symmetric nearest neighbors in 3D. This process was performed in each nuclear ROIs. Point combinations that constructed only a single barcode were immediately matched to the on-target barcode set. 2 rounds of error corrections were implemented out of 5 total barcoding rounds. For points that matched to multiple barcodes, the point sets were filtered by calculating the residual spatial distance of each potential barcode point set and only the point sets giving the minimum residuals were used to match to a barcode. If multiple barcodes were still possible, the point was matched to its closest on-target barcode with a hamming distance of 1. If multiple on target barcodes were still possible, then the point was dropped from the analysis as an ambiguous barcode. This procedure was repeated using each hybridization as a seed for barcode finding and only barcodes that were called similarly in at least 4 out of 5 seeds were used in the analysis. For the false positive estimates, both blank barcodes and on-target barcodes were run simultaneously. Those blank barcodes consisted of all the remaining barcodes out of 2,048 barcodes that allow 2 rounds of error corrections in 5 total barcoding rounds.

For channel 3 decoding in DNA seqFISH+, once all potential points in the first 60 hybridizations (hyb 1-60) were obtained, intensities of all the potential chromosome paint partners in the other 20 hybridizations (hyb 61-80) were computed on the pixels where points were found. At this step, each point has 20 intensity values, corresponding to those from individual chromosome paints. Those chromosome paint intensities found on the points in nuclei from all positions and all hybridization rounds (hyb 1-60) were grouped by chromosome, and then z score was calculated. The z score values were thresholded with 1, and each point was assigned with unique chromosome identity, whose value was above the threshold. Only a minimum fraction of points (<3%) were assigned to multiple chromosomes and dropped as ambiguous points.

For the sequential immunofluorescence image processing, in contrast to spot detection processing as described above, background subtraction was not applied to the images, except for RNAP II Ser5-P visualization shown in Fig. 3d and Extended Data Fig. 7d. The alignment and correction for chromatic aberration shifts between different fluorescent channels were performed as described above. Then intensity values for all the voxels within individual nuclear ROIs were obtained for all IF channels as well as repetitive elements (telomere, MinSat, LINE1 and SINEB1) and DAPI.

### Analysis of sequencing-based data

Hi-C data was obtained from NCBI GEO (accession GSE96107) and processed as described before^10^. SPRITE data were obtained from the 4D Nucleome data portal (data.4dnucleome.org, accession 4DNESOJRTZZR). ChIP-seq data were obtained from ENCODE (encodeproject.org) as bigWig tracks and the average relative signal in each genomic bin was calculated using the UCSC Genome Browser program bigWigAverageOverBed. DamID data were obtained from NCBI GEO (accession GSE17051) and the genomic coordinates of DamID microarray probes were converted from mm9 to mm10 using the UCSC Genome Browser program liftover. DamID values were calculated as the mean DamID score within each genomic bin. Repli-seq data were obtained from NCBI GEO (accession GSE102076) and the replication timing at each genomic bin was calculated as the log2 ratio of early and late S fractions. GRO-seq data were obtained from NCBI GEO (GSE48895) and aligned to mm10 using bowtie2 to create bam files. Read counts at each genomic bin were obtained from bam files using bedtools multicov. Hi-C data was binned at the 25 kb and 1 Mb resolution, and all the other data were binned at the 1 Mb resolution.

### Visualization of seqFISH data

DNA seqFISH+ data were visualized using Pymol 2 by generating a .xyz file containing the x,y,z coordinates of each FISH probe coordinate. Each coordinate was displayed as a sphere, and sticks were drawn between coordinates that were consecutive in the genome. Immunofluorescence signal was visualized by displaying a surface around x,y,z coordinates with intensity Z-score values above 3.

### DNA contact map analysis

To generate a pairwise contact map from the DNA seqFISH+ dataset, for each locus in a single cell, the identities of other loci within a search radius of 500 nm for channel 1 and 2 and 150 nm for channel 3 were tabulated. The total occurrence of any pairwise interaction was normalized by the product of the occurrence frequency of each of the loci. The contact map was compared with the Hi-C contact map^4^ in Figure 1. The contact maps for all chromosomes for both 1 Mb and 25 kb data are shown in Extended Data Fig. 3.

### Physical distance vs genomic distance

In each cell, two sister chromosomes were separated by finding the consensus between two clustering algorithms: Spectral method in the FindClusters function in Mathematica and Ward method. For most chromosomes in single cells, the two copies of sister chromosomes occupied distinct regions in the nucleus, while in some cells, they were fused together. In a small percentage of cells, 3 or more alleles of the same chromosome could be observed. However, in a vast majority of cells, only 2 chromosomal territories were observed indicating that replicated chromosomes mostly stay together^67^ until segregation. For the 25 kb data, the alleles were separated by the DBSCAN clustering algorithm in scikit-learn library in python.

Along each allele of a given chromosome in single cells, we calculated the physical distances between all pairs of detected loci and paired them with their genomic distances. For a fixed genomic distance, the median physical separation values are shown in Fig. 1i for the 1 Mb data for all the chromosomes, and Fig. 1j for the 25 kb resolution data.

The same genomic vs physical distance scaling was calculated for loci belonging to the same zone assignments. We calculated a “return probability,” i.e. the probability for a locus to return to within 500 nm of the reference locus either when both loci were in the same zone or were just on the same chromosome. The return probability from all chromosomes are then combined based on the genomic distances. The return probability for two zones are shown in Fig. 2g as well as the return probability for all loci along a chromosome. The return probabilities for all other zones and interfaces are shown in Extended Data Fig. 8a.

### IF normalization and clustering analysis

For the voxel-based multiplexed IF analysis, we first aligned the sequential immunofluorescence data across all rounds of hybridization (see ‘Image Analysis’). Then voxels in each channel were binned 2×2×1 (200 nm x 200 nm x 250 nm), because the diffraction limit is approximately 200-250 nm in the fluorescence channels imaged. All subsequent data analysis were performed on the binned data. Because tens of millions of voxels from all of the cells were too numerous for clustering analysis, a representative subset of voxels are selected, clustered and used as a training set to train a model which then propagated the cluster identification to all voxels in the data. To do so, voxels from a single Z plane (plane 13, approximately midpoint in the cell) out of 25 z-slices for all cells were selected. In each cell, individual channels were z-score normalized. The voxels with total z-score values more than 0 summed over 16 IF channels were selected and normalized by the total z-score to account for voxel to voxel intensity variations. All pixels of the cells within the first experiment (n = 201 cells) were then combined and one out of every 200 pixels are selected and clustered by hierarchical clustering using the Mathematica Agglomerate function and Ward distance option. 10 clusters or nuclear zones were assigned to all 60,482 pixels as the training set. These classified zone definitions were then propagated to the rest of the pixels in each cell normalized by the above procedure using the GradientBoostedTree option in the Classify function in Mathematica. Separately, pixels with lamin and Fibrillarin marker z-score >1 were assigned to the nuclear lamin and nucleolus zones. The 44,000 pixels, which are assigned to one of the 12 nuclear zones and contain 16 intensity values from individual IF markers, were then visualized in Fig. 3b with Uniform Manifold Approximation and Projection (UMAP)^42^ using a umap-learn library in python.

### Association of loci with zones

For each DNA loci decoded from the DNA seqFISH+ experiment, the nearest pixels within 300 nm and the zone assignments for those pixels were collected in each cell. It is possible to have a locus be in contact with multiple zones. If a locus interacts with more than two zones, for example Pou5f1 (Oct4) in cell 38 is interacting with zone 1, 2, and 8, then its zone interactions were divided into pairs of zones, or “interfaces”. In other words, that locus was counted 1/3 toward each of the interfaces (1, 2), (2, 8), and (1, 8). For individual loci, the frequencies of appearing in all zones and interfaces were normalized to unity and shown in Fig. 2c. For the analysis shown in Fig. 3e, the total number of DNA loci detected each zone and interfaces are tabulated and normalized to unity for each zone or interfaces between pairs of zones. The same analysis for zone proximity was performed on the set of loci that are interacting with other loci on the same chromosomes (intrachromosomal) and with loci on the other chromosomes (interchromosomal) within 300 nm. Similarly, the introns from the 1,000 gene experiments were tabulated for their zone and interface assignments. Randomized DNA loci were generated by selecting a random set of voxels in the nucleus while keeping the total number of DNA loci the same in a given cell. Then the voxels were offset by a random xyz value with a 100 nm radius. To bootstrap all of the data sets, we randomly sampled 150 cells out of n = 201 cells with 20 trials and calculated the mean and standard errors.

For each loci, a set of zone and interface occupancy were tabulated and shown in Fig. 3f. Autocorrelation function for each zone was calculated along the chromosome for all the loci to measure the “persistence length” of loci associated with each zone and is shown in Extended Data Fig. 8b.

To determine the “Chromatin profile” for each loci shown in Fig. 2c, the nearest DNA loci within 300 nm of each antibody voxel that has a normalized z-score above 2 are collected. Similar analysis is applied to DAPI and the repetitive element data. The results are tabulated and totaled across all cells (N=201) for each loci. These results are compared with ChIP-seq^8^, SPRITE^10^ and DamID^7^ datasets in Fig. 2d and Extended Data Fig 5d.

The chromatin profiles for all loci were clustered by hierarchical clustering using the Agglomerate function in Mathematica with the Ward distance option and plotted in tSNE with scikit-learn library in python in Fig. 2e. 15 chromatin marks along with DAPI were used, and 4 clusters were selected. Cluster 1 is enriched in repressive markers such as H3K9me3, mH2A1 and DAPI. Cluster 2 was enriched in interactions with Fibrillarin and associated with nucleolus. Cluster 3 was enriched in active marks such as RNAP II Ser5-P, H3k27ac and SF3a66 (speckle marker). Cluster 4 was enriched in Lamin B1. In individual cells, loci associated with each cluster were mapped onto the chromosome structure images shown in Fig. 2g. To calculate the interactions within and between clusters, we computed the frequency of finding a loci from a given cluster in contact within a 1 μm radius with another loci of the same or different cluster identity. The total number of intra-cluster and inter-cluster interactions were tabulated and normalized to unity. Randomized data was generated by scrambling the cluster identities of individual loci in cells while keeping the total number of loci within each cluster the same within that cell. The interaction frequency for observed and randomized data for each cell are shown as boxplots in Fig. 2f. Similar analysis is performed for A/B compartment assignments^4^, and shown in Extended Data Fig. 6d-g. The loci without A/B compartment assignments in the study were excluded from the analysis.

### Correlation of zone with gene expression

To calculate the correlation between expression and zone assignment, we took each channel 1 and 2 locus and computed the total RNAseq TPM values^34^ within 50 kb upstream and downstream of that locus. We normalized the total frequency of appearing in one of the zones or interfaces to unity for each loci. We then correlated the Log (1+expression value) of all 2,460 regions with the frequency of finding them in each of the zones/interfaces. Similar analysis was performed for GRO-seq^46^ using Log (1+GRO-seq value) and Replication timing^47^ datasets with mESCs.

To determine whether we can predict the mean expression values for each locus based on its zone association profiles, we estimated the expression level for a given loci as a sum of the product between the normalized frequency of being in each zone/interface for that loci and the Pearson correlation coefficient between the zone/interface with the mean expression value across all the loci. The estimated expression values for all 2,460 loci were correlated with the actual expression values with a Pierson’s coefficient of 0.54.

For calculating the correlation between mRNA expression levels with zone assignments in single cells, we first z-scored the single cell mRNA seqFISH measurements for 27 genes after normalizing by Eef2 expression levels to account for cell size differences and selecting cells in the center field of view (n = 125). Then we counted the frequency of each of the measured loci contacting a voxel with an active or speckle zone assignment (zone 1 and 2) within 300 nm, normalized by the total number of voxels that loci were in contact within 300 nm. The Pearson coefficient was computed between the z-scored expression value and the active/speckle zone contact frequency. To randomize the sample, we shuffled the z-score normalized expression values with active/speckle zone occupancy from different cells over 20 randomized trials. The correlation coefficient for each gene was calculated and plotted in Fig. 3i.

For calculating the correlation between intron expression levels with zone assignments in single cells, we classified the corresponding DNA loci as “ON” or “OFF” based on whether introns were bursting at that loci or not for all 14 introns measured. Then the active and speckle zone occupancy for loci in each category was calculated and shown in Fig. 3j with each point representing one intron.

### Colony analysis

For cells within the unlabeled E14 colonies, we compute the correlation of the IF states, RNA states and chromosome structures between pairs of cells. Individual RNA levels were normalized by Eef2 expression level and then z-scored across all cells in the experiment. The chromosome contact correlations between cells are computed as follows. First, a 20×20 chromosome to chromosome contact matrix is generated for each cell with a search radius of 2.5 μm. Then the correlations between cells were computed as the Pearson coefficient of the entries of the two matrices. The intensity of individual IF marks was first normalized by the total intensity of all IF marks and then z-scored within each field of view. The averages of the cell pair correlation values for IF, RNA and chromosomes are shown in Fig. 4f for 24 hour and 48 hour clonal tracing as well as controls (correlation of pairs of cells between colonies in the 24 hour and 48 hour data). In addition, we computed the variance of individual IF marks within single colonies in the 48 hour experiment compared to the variance between cells of different colonies. IF marks that have longer memories showed lower variance within colonies compared to the variance between colonies in Fig. 4h.

### Normalization of global chromatin levels in single cells

To remove the contributions from cell size, background signals, the affinity of antibody used, as well as differences between replicates, we constructed a generalized linear model (GLM) for the sequential immunofluorescence data using the glm() function in R, which had been used to adjust for systematic bias in single cell RNA sequencing data^68–70^, for each chromatin mark *i*, using a Gaussian error distribution:

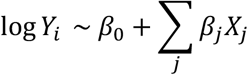

*Y_i_* represents the vector of total fluorescence intensity of chromatin mark *i* across all cells, and *X_j_* is a vector of latent variables contributing to the systematic bias in global chromatin states quantification. We included cell size (*μm*^2^), total fluorescence intensity over all chromatin marks per cell, experimental replicate ID and field of view (FOV) ID as latent variables in the GLM, and used the Pearson residuals of each fitting as the corrected standardized values of single cell chromatin state.

### Characterizing the heterogeneity of global chromatin states in single cells

Using the adjusted global chromatin states quantification, we then aimed to identify heterogeneity across single cells using clustering approach. In contrast from single cell RNA-seq data where single cells are represented by high-dimensional gene expression vectors, global chromatin state is of lower dimension and dimensionality reduction approach such as principal component analysis (PCA) is not necessarily required. We computed the pairwise Euclidean distance of single cells, which was used to compute a K-nearest neighbor (KNN) graph with K = 10. The KNN graph was then used as the input for Uniform Manifold Approximation and Projection (UMAP)^42,71^ as two-dimensional visualization, and was also subsequently transformed into a shared nearest neighbor (SNN) graph for Leiden clustering^72^. Distance matrix was computed using the R function dist(). The distance matrix was used for KNN and SNN graphs-finding using the FindNeighbors() function, and UMAP was computed on the KNN matrix using RunUMAP() (calling the umap-learn Python function) in the R package Seurat. Leiden clustering is performed on the SNN graph using the R Seurat function FindClusters() with the argument algorithm = 4. To explore cell-cell variation in the global chromatin states other than cell cycle phases, the four cell cycle markers (Geminin, H4K20me1, H3pSer10 and H4K16ac) were removed from KNN graph construction and the process of UMAP and Leiden clustering in Fig. 4b, c. Then we mapped back the expression levels of the four cell cycle markers onto the clustering results in Extended Data Fig. 9d, confirming that the cell clusters obtained by immunofluorescence data were not affected by cell cycle phases.

We then examined if a systematic quantification of cell cycle could be achieved with global chromatin states, and correspondingly constructed a principal curve^73^ with principal_curve() function from the R package princurve, using the four cell cycle markers from the immunofluorescence data – Geminin, H4K20me1, H3pSer10 and H4K16ac. We found that the single cell projection onto the resulting principal curve depicted a progression from G2/M to S phase, due to H4K20me1 (a marker for G2/M phases) and H4K16ac (a marker for S phase)^74^ accounting for most of the variance in single cell data. We also confirmed that the majority of other chromatin markers were not affected by cell cycle phases, by projecting single cell data onto the principal curve for individual chromatin marks, shown in Extended Data Fig. 9e.

### Characterizing transcriptional heterogeneity of single cells

Similar to global chromatin states quantification, we constructed a GLM for individual gene expression vector in RNA data, with cell size, total profiled transcripts per cell, experimental replicate ID and FOV ID as latent variables. Pearson residuals were taken as the corrected and standardized expression values, and were subsequently used for computing the KNN graph with K = 10, and the KNN graph was used as the input for UMAP visualization and Leiden clustering in Extended Data Fig. 9a, b. IF marks profiles are shown in the RNA clusters as well.

### Reporting summary

Further information on research design is available in the Nature Research Reporting Summary linked to this paper.

### Code availability

The custom written scripts used in this study are available at https://github.com/CaiGroup/dna-seqfish-plus.

### Data availability

Source data from this study are available at figshare (DOI: 10.6084/m9.figshare.12089412). All raw data obtained during this study are available from the corresponding author upon reasonable request.

